# Quantifying Data Distortion in Bar Graphs in Biological Research

**DOI:** 10.1101/2024.09.20.609464

**Authors:** Teng-Jui Lin, Markita P. Landry

## Abstract

Over 88% of biological research articles use bar graphs, of which 29% have undocumented data distortion mistakes that over- or under-state findings. We developed a framework to quantify data distortion and analyzed bar graphs published across 3387 articles in 15 journals, finding consistent data distortions across journals and common biological data types. To reduce bar graph-induced data distortion, we propose recommendations to improve data visualization literacy and guidelines for effective data visualization.

## Main

Bar graphs are widely used in biological research because relative changes between experimental groups are easily compared visually^1,2^. Therefore, correctly creating and interpreting bar graphs is crucial for understanding the underlying data. To accurately represent data with the bar’s area, bar graphs need to follow the “principle of proportional ink” (Ref ^3^)—the area of each bar needs to be directly proportional to the value being represented. Violation of this principle causes data distortion in area-based visualizations. For example, not starting bars at a zero y-axis value (zeroing mistakes) distorts the underlying data^1,3–6^, often by overrepresenting findings (Extended Data Fig. 1).

However, we noted a more nuanced distortion in bar graphs that has been overlooked in research: the distortion caused by using logarithmic axes (log mistakes). Log mistakes occur when areas of the graph (perceived directly proportional to their value) are used to represent the much larger scale difference of log-spaced values, which can drastically change the immediate interpretation of the graph. Depending on the numerical variables represented by the graph, called measurands (e.g. ratio, relative bioluminescence, relative gene expression, or cytokine concentration), and measurand types (e.g. absolute/relative), accurate representation of log-spaced values requires proper data transformation, axis scaling, and even alternative graph types, as demonstrated by examples (Extended Data Fig. 2–5).

Despite the common use of bar graphs in biological research, the prevalence of their use and distortion remains underexplored. The extent of data distortion caused by the two main types of bar graph mistakes—zeroing and log mistakes—is also not fully understood, presumably due to a lack of rigorously characterized metrics. Here, we developed a mathematical framework to quantify the extent of data distortion caused by zeroing and log mistakes (Supplementary Discussion) and applied the framework to bar graphs published across 3387 articles in 15 high-impact journals in 2023 (Methods). We suggest potential causes of bar graph-based data distortion and propose recommendations for improving data visualization literacy and standardizing bar graph usage across biological research.

We first investigated the prevalence of bar graph visualization mistakes that violate the principle of proportional ink (Methods). Over 88% of 3387 sampled biological research articles contain at least one bar graph (Fig. 1a), consistent with previous findings in physiology journals^7^. Of the 2985 articles with bar graphs, 29% contain at least one visualization mistake (Fig. 1b). This striking trend is consistent across journals regardless of publisher or presence of an art editor. Of the 874 articles with visualization mistakes, 60% have zeroing mistakes, 40% have log mistakes, and 12% have other mistakes (Fig. 1c), suggesting that both zeroing and log mistakes have been overlooked. Interestingly, articles with bar graphs, especially those with visualization mistakes, have more authors than those without (Extended Data Fig. 6), consistent with our hypothesis that having more authors increases the probability of having coauthors who contribute visualization mistakes.

**Fig. 1.**
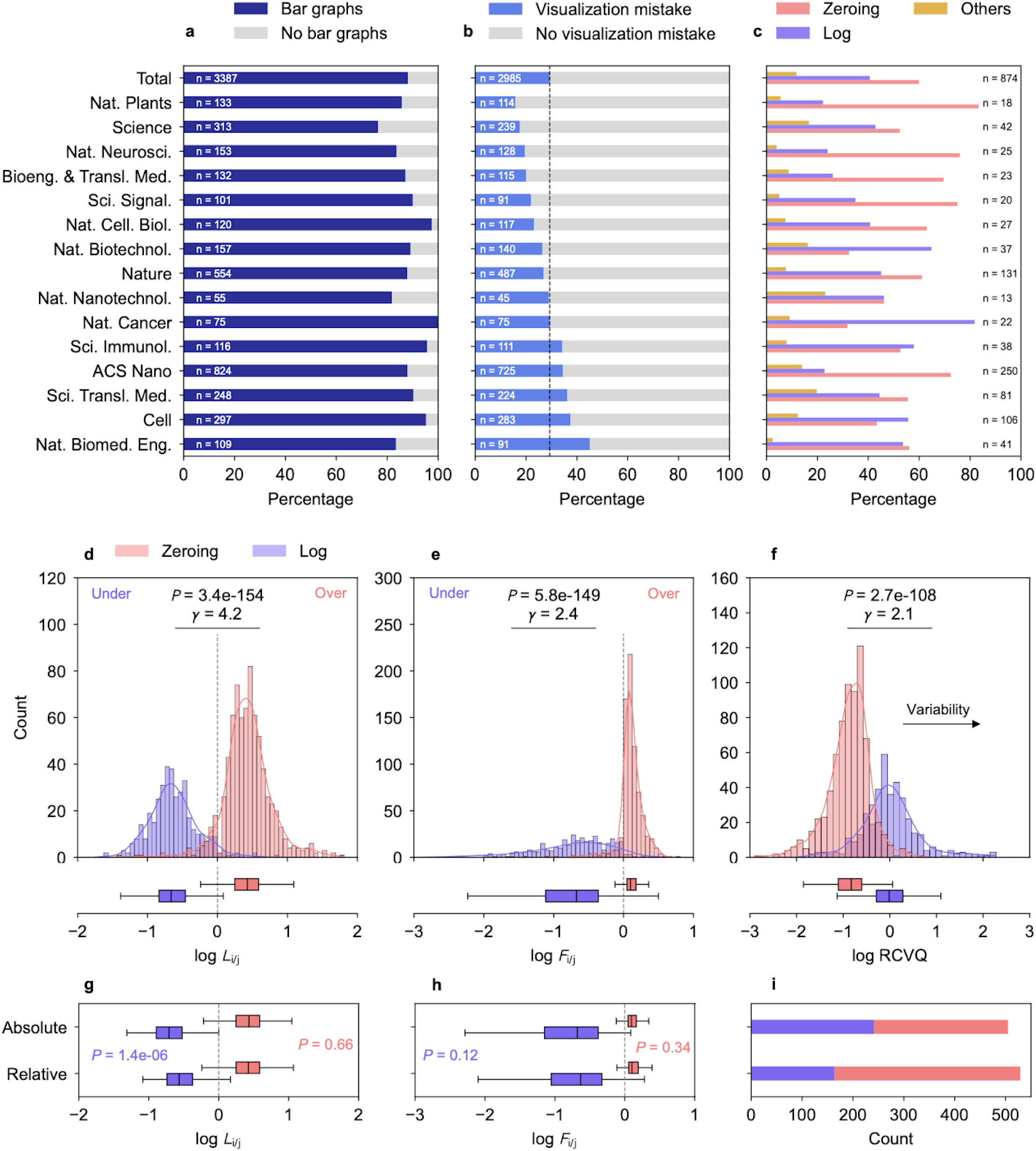
Prevalence and extent of data distortion in bar graphs in biological research. **a**, Percentage of articles with or without bar graphs in *n* articles retrieved from each journal. **b**, Percentage of articles with or without bar graph visualization mistakes in *n* articles with at least one bar graph. Dashed line represents the percentage of articles with visualization mistakes from all sampled articles. **c**, Percentage of articles with zeroing (pink), log (purple), and other (gold) bar graph visualization mistakes in *n* articles with bar graph visualization mistakes. **d, e**, Histograms and box plots of bias-mitigated log-transformed lie factor of relative change *L*_*i/j*_ (**d**) and fold change *F*_*i/j*_ (**e**) for zeroing (*n* = 747) and log (*n* = 387) mistakes. Histogram bins outside the displayed domain (<0.7% of the population) are not plotted for clarity. Gray dashed lines represent metric for the correct visual representation. **f**, Histograms and box plots of bias-mitigated log-transformed RCVQ of mark proportionality constants for zeroing (*n* = 732) and log (*n* = 390) mistakes. Histogram bins outside the displayed domain (<0.8% of the population) are not plotted for clarity. Colored solid lines represent kernel density estimations. *P* values comparing median of zeroing and log groups were computed using two-sided Mann-Whitney U test. **g, h**, Box plots of bias-mitigated log-transformed lie factor of relative change (**g**) and fold change (**h**) for representations of absolute or relative measurands hued by zeroing and log mistakes. Gray dashed lines represent well-representation. *P* values comparing median of absolute and relative groups for zeroing (pink) or log (purple) mistakes were computed using two-sided Mann-Whitney U test. **i**, Number of graphs zeroing and log mistakes in representations of absolute or relative measurands. Box plot shows 25, 50, and 75 percentiles. Whiskers extend to the farthest data point within 1.5 times the interquartile range. Effect sizes were computed using nonparametric gamma. RCVQ, robust coefficient of variation based on interquartile range.

We then quantified the extent of data distortion to understand how zeroing and log mistake types cause misrepresentation of data’s relative change [(*x*_2_ – *x*_1_) / *x*_1_] and fold change (*x*_2_ / *x*_1_) between experimental groups (Methods). We found that zeroing mistakes significantly overrepresent the data’s relative and fold changes by 2.7 and 1.3 times, whereas log mistakes significantly underrepresent the data’s relative and fold changes by 4.6 and 4.9 times, respectively (Fig. 1d, e). The magnitude of distortion of both relative and fold change is larger for log mistakes than zeroing mistakes, suggesting log mistakes are more visually misleading than zeroing mistakes. This is especially concerning because using logarithmic axes in bar graphs is currently a standard practice in the field. Furthermore, the magnitude of data distortion for fold changes is larger than that of relative changes for both zeroing and log mistakes (Extended Data Fig. 7), suggesting the audience is more misled when interpreting fold change than relative change. The larger distortions in log mistakes over zeroing mistakes are further supported by their larger variability of mark proportionality constants (Fig. 1f), the ratio between true and visualized values that should be the same for each bar, indicating log mistakes’ larger deviation from the principle of proportional ink.

We further explored the relationship between data distortion and measurand types (i.e. absolute/relative), measurand identity (e.g. transcript levels, relative fluorescence), and journals. First, we found no difference between data distortion in absolute (e.g. concentration) and relative (e.g. ratio) measurands (Fig. 1g, h), though we note that the distortion of relative change for log mistakes is larger for absolute measurands than for relative measurands (Fig. 1g). Furthermore, while the number of misused bar graphs representing absolute and relative measurands is similar, graphs representing absolute measurands are equally likely to make zeroing and log mistakes, whereas those representing relative measurands are more likely to make zeroing mistakes (Fig. 1i). As a result, relative measurands—often used to represent percentages and ratios—tend to overrepresent data’s relative and fold changes.

Among all measurands represented by misused bar graphs, percentage, ratio, concentration, and count are the most common (Extended Data Fig. 8a–e). Zeroing mistakes are more common for percentage measurands, whereas log mistakes are more common for count measurands. Zeroing and log mistakes are equally common for concentration and ratio measurands. Temperature measurands are only observed in zeroing mistakes and have the most relative change distortion. Note that temperature on a relative scale (e.g. Celsius and Fahrenheit scales instead of Kelvin) should not be plotted using bar graphs for relative change comparisons because relative changes are ill-defined for measurands with arbitrarily defined zero baselines (e.g. 2 °C is not twofold of 1 °C). Instead, mean-and-error plots are better suited for measurands with undefined relative changes.

Among the 15 high-impact journals sampled, *Nature Biomedical Engineering, Cell, Science Translational Medicine*, and *ACS Nano* have the highest percentage of articles with visualization mistakes (Fig. 1b). *Nature* and these journals (except *Nature Biomedical Engineering*) have the highest number of misused bar graphs and similar levels of data distortion among themselves (Extended Data Fig. 8f–i). While *ACS Nano* articles mostly make zeroing mistakes, *Nature, Nature Biomedical Engineering, Science Translational Medicine*, and *Cell* articles are equally prone to zeroing and log mistakes (Extended Data Fig. 8j).

The prevalence of misused bar graphs is potentially caused by a systematic lack of data science training in science, technology, engineering, and mathematics (STEM) education, research training, and the academic publication cycle (Fig. 2a). At all levels of STEM education, data science courses remain optional for most programs despite their importance in research and in the industrial workforce^8–10^. In the publication cycle, responsible personnel—from first authors and corresponding authors to peer-reviewers and editors—frequently overlook errors in data visualizations (Fig. 1b), suggesting the need for continued data science training.

**Fig. 2.**
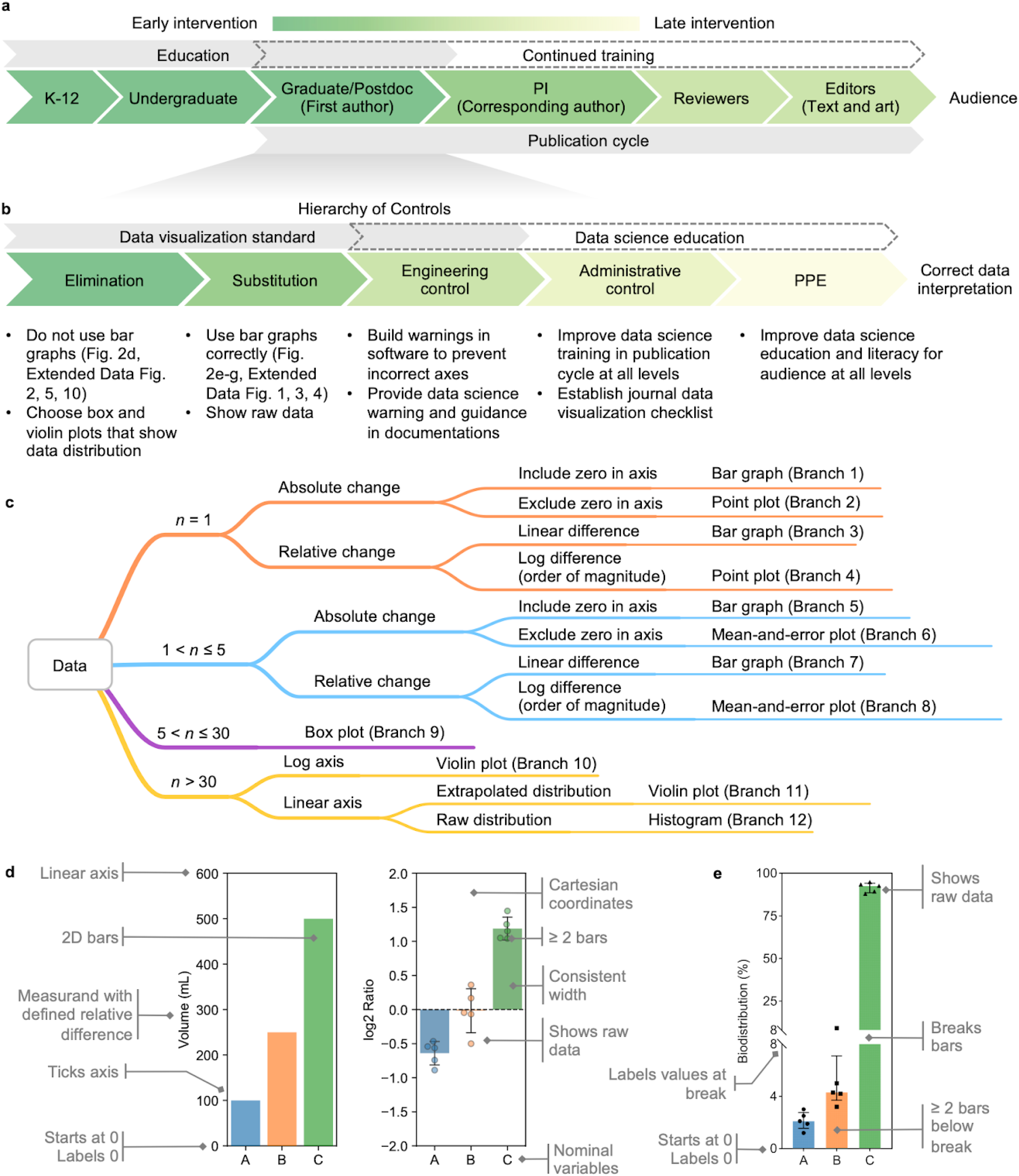
Recommendations to prevent bar graph misuse and improve data visualizations for comparing numerical values between nominal groups. **a**, Systematic lack of data science training exists in education, research training, and the publication cycle. K-12, from kindergarten to twelfth grade; PI, principal investigator. **b**, Hierarchy of controls^11^ applied on an individual level decreases the incidence of bar graph misuse. Colormap denotes the timeliness of interventions for preventing the publication of misused bar graphs and facilitating correct data interpretation. PPE, personal protective equipment. **c**, Decision tree for choosing graph types (Extended Data Fig. 10). **d, e**, Guidelines for making bar graphs, including general guidelines (**d**) and additional guidelines for bar graphs with discontinuous axes (**e**).

To prevent bar graph misuse, we took inspiration from the hierarchy of controls^11^ for laboratory hazards and developed intervention strategies for personnel on all levels to minimize data misinterpretation (Fig 2b). As the earliest intervention, eliminating bar graphs and using alternative graph types will allow clearer visualization of data distribution and avoid data distortion. We created a decision tree for selecting appropriate graph types (Fig 2c, Extended Data Fig. 9) and stress that the goal of data visualization is to convey scientific messages accurately and accessibly, but not to make eye-catching figures. For example, bar graphs in polar coordinates, such as circular and radial bar graphs, inherently distort the underlying data and should not be used (Extended Data Fig. 10, Supplementary Discussion). When the decision tree indicates that bar graphs are necessary for showing relative changes, visualization guidelines need to be adhered to avoid data distortion. We developed guidelines for making bar graphs, including those with discontinuous (broken) axes (Fig. 2d, e). When bar graphs are being made, these guidelines should also be enforced on a software level by implementing engineering controls, such as built-in warnings and tutorials, which are currently lacking (Supplementary Discussion). When bar graphs are being misused, administrative controls like journal editorial checklists and reviewer oversight can prevent their publication. We propose data visualization checklists for authors, reviewers, and editors (Supplementary Discussion) and suggest providing regular journal personnel training for an improved understanding of data visualization standards, especially at the journal editor level. As the final layer of protection against incorrect data interpretation, improving data science literacy by expanding data science education coverage will empower the audience to identify data misrepresentations and interpret results accordingly.

In conclusion, we found that an alarming 29% of articles with bar graphs contain misuses despite their widespread use in biological research. We developed data distortion metrics and found zeroing mistakes distort data’s relative and fold change by 2.7 and 1.3 times, usually by overrepresentation. Conversely, log mistakes distort data’s relative and fold change by 4.6 and 4.9 times, usually by underrepresentation. These mistakes and distortions might significantly impact researcher’s interpretation of biological data. We suggest inadequate data science education as a key factor behind these issues and recommend implementing guidelines for effective data visualization along with strategies to systematically reduce data misrepresentation.

## Methods

### Bar Graph Categorization

PDFs of research articles published online in 2023 in high-impact journals in biological and biomedical sciences and engineering were downloaded using search queries from the publisher’s websites (Supplementary Methods). The downloads are spread out into multiple sessions in small amounts to avoid systematic downloading. Other article types, such as reviews, article corrections, article retractions, retracted articles, matters arising articles, and response articles, are excluded. All figures presented in the PDFs are included for categorization. Bar graphs are broadly defined as graphs that use bars or lines as marks and their area and length as magnitude channels, such as conventional bar graphs, histograms, and lollipop graphs. Each article is categorized into three mutually exclusive categories: (1) contains no bar graphs, (2) contains at least 1 bar graph and all the bar graphs do not have visualization mistakes, and (3) contains at least 1 bar graph and at least 1 of the bar graphs have visualization mistakes. Visualization mistakes of bar graphs are defined and categorized into three mutually exclusive categories in order: bar graph with (1) a linear y-axis and nonzero bar baseline (zeroing mistake), (2) a logarithmic y-axis (log-mistake), and (3) other mistakes. Other mistakes consist of bar graphs (1) in polar coordinates, (2) in nonstandard coordinates (e.g. logarithmic axis with zero or negative values), (3) with inconsistent bar width, (4) with three-dimensional (3D) bars, (5) with overlapping bars, (6) with breaks that do not have ≥ 2 bars under the break, (7) with bars of non-rectangular shapes. Articles with incorrectly visualized bar graphs are assigned with all the applicable non-mutually exclusive annotations of zeroing, log, and other mistakes.

### Quantification of Bar Values

All misused bar graphs are screenshotted individually. Graphs with shared y-axis tick labels are considered one graph. The bar graphs with zeroing and log mistakes are digitized by WebPlotDigitizer v5 (Ref ^12^) (https://automeris.io/). For each bar graph, the axis is first calibrated to its true value using the axis ticks, and the value corresponding to each bar is recorded as the true value. The axis is then calibrated from the baseline of the bars to the topmost axis tick mark using a linear scale from 0 to 1. The value corresponding to each bar is then recorded as the visualized value. Unquantifiable bars are excluded from digitization if they (1) have zero height, (2) have breaks, or (3) have unidentifiable top edge due to the overlay of raw data points. All bars from a graph are excluded if the graph is (1) a histogram, (2) a stacked bar graph, (3) a lollipop graph, (4) in polar coordinates, (5) in nonstandard coordinates, (6) consisted of only one quantifiable bar, (7) consisted of >30 quantifiable bars, (8) consisted of 3D bars, or (9) not ticked with y-ticks.

### Measurand Annotation

The y-axis labels for bar graphs with zeroing and log mistakes are manually recorded as the measurand represented by the bar graph. The measurand annotations are further categorized within four successively narrower scope: (1) absolute or relative measurands, (2) measurement type, (3) manually-curated detailed measurand annotations with a broader scope (Measurand Level II), and (4) manually-curated measurand annotations with a narrower scope (Measurand Level I) (Supplementary Methods). The categories within each scope are mutually exclusive.

### Data Distortion Metrics

Well-represented graphs are graphs that follow the principle of proportional ink. Well-behaved graphs are graphs that do not represent true values with visualized values of opposite signs. Let each bar’s true value be *x*_*i*_ and visualized value be *y*_*i*_. We developed and defined mark proportionality constant as

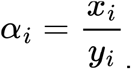

The lie factor of relative change between bars *i* and *j* is defined as^3^

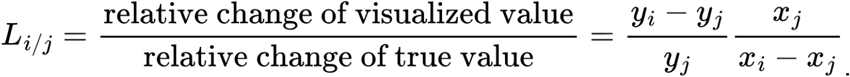

For well-behaved graphs, the lie factor of relative change quantifies distortion of relative changes on a pair-wise permutation level (Supplementary Discussion):

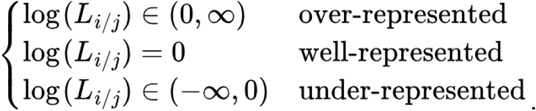

We developed the lie factor of fold change between bars *i* and *j* and defined it as

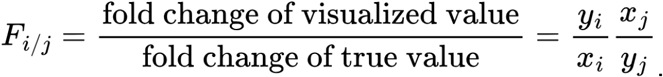

For well-behaved graphs, the lie factor of fold change quantifies distortion of absolute changes on a pair-wise combination level (Supplementary Discussion):

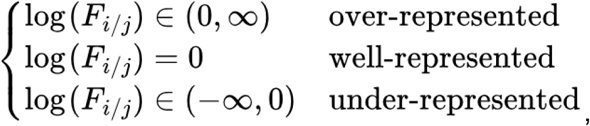

for *i*-*j* combinations such that |*x*_*i*_| > |*x*_*j*_|.

We took inspiration from statistics to find metrics to quantify the variability of mark proportionality constants. The normalized variability metrics are based on interquartile range (IQR) or median absolute deviation (MAD)^13^. The robust coefficient of variation based on IQR (RCVQ) is defined as

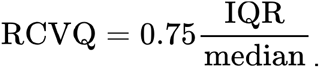

The robust coefficient of variation based on MAD (RCVM) is defined as

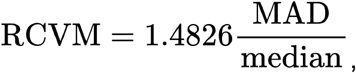

where MAD is defined as the median of the set of absolute deviations of each element in the population with the median of the population. The RCVQ and RCVM of mark proportionality constants are zero for well-represented graphs and positive for misrepresented graphs.

The mathematical framework of data distortion metrics, including detailed derivation, theoretical characterization, and discussion, is presented in Supplementary Discussion.

### Statistics

Bar-level bias is mitigated by taking the median of bar-level metrics within each graph (Supplementary Discussion). Graph-level bias is mitigated by grouping graphs within each article by mistake type and Measurand Level I, and taking the median of the metric within each grouped set of graphs.

*P* values for comparing between two groups are calculated with two-sided Mann-Whitney U test. *P* values for comparing one group with a baseline of zero are calculated with two-sided quantile test (second quantile, median). Effect sizes are calculated with nonparametric Cohen’s *d*-consistent effect size *γ* (Supplementary Methods)^14^.

## Declarations

### Data Availability

All data presented in this article, including article metadata, screenshots of misused bar graphs, value annotations, and measurand annotations are available at https://github.com/tengjuilin/misused-bar-graphs.

### Code Availability

All code used for data analysis and visualization is also available at https://github.com/tengjuilin/misused-bar-graphs.

## Acknowledgments

We acknowledge support of a Burroughs Wellcome Fund Career Award at the Scientific Interface (CASI) (MPL), a Dreyfus foundation award (MPL), the Philomathia foundation (MPL), an NSF CAREER award 2046159 (MPL), an NSF CBET award 1733575 (MPL), a CZI imaging award (MPL), a Sloan Foundation Award (MPL), a McKnight Foundation award (MPL), a Simons Foundation Award (MPL), a Moore Foundation Award (MPL), a Brain Foundation Award (MPL), and a Schmidt Science Polymath Award from Schmidt Sciences, LLC (MPL). MPL is a Chan Zuckerberg Biohub investigator and a Hellen Wills Neuroscience Institute investigator.

## Author Contributions

Conceptualization: T.-J.L.; methodology and experimental design: T.-J.L.; data collection, analysis, and visualization: T.-J.L.; writing (original draft): T.-J.L.; writing (review and editing): T.-J.L., M.P.L.; project supervision: M.P.L.; funding acquisition: M.P.L.

## Figures

**Extended Data Fig. 1.**
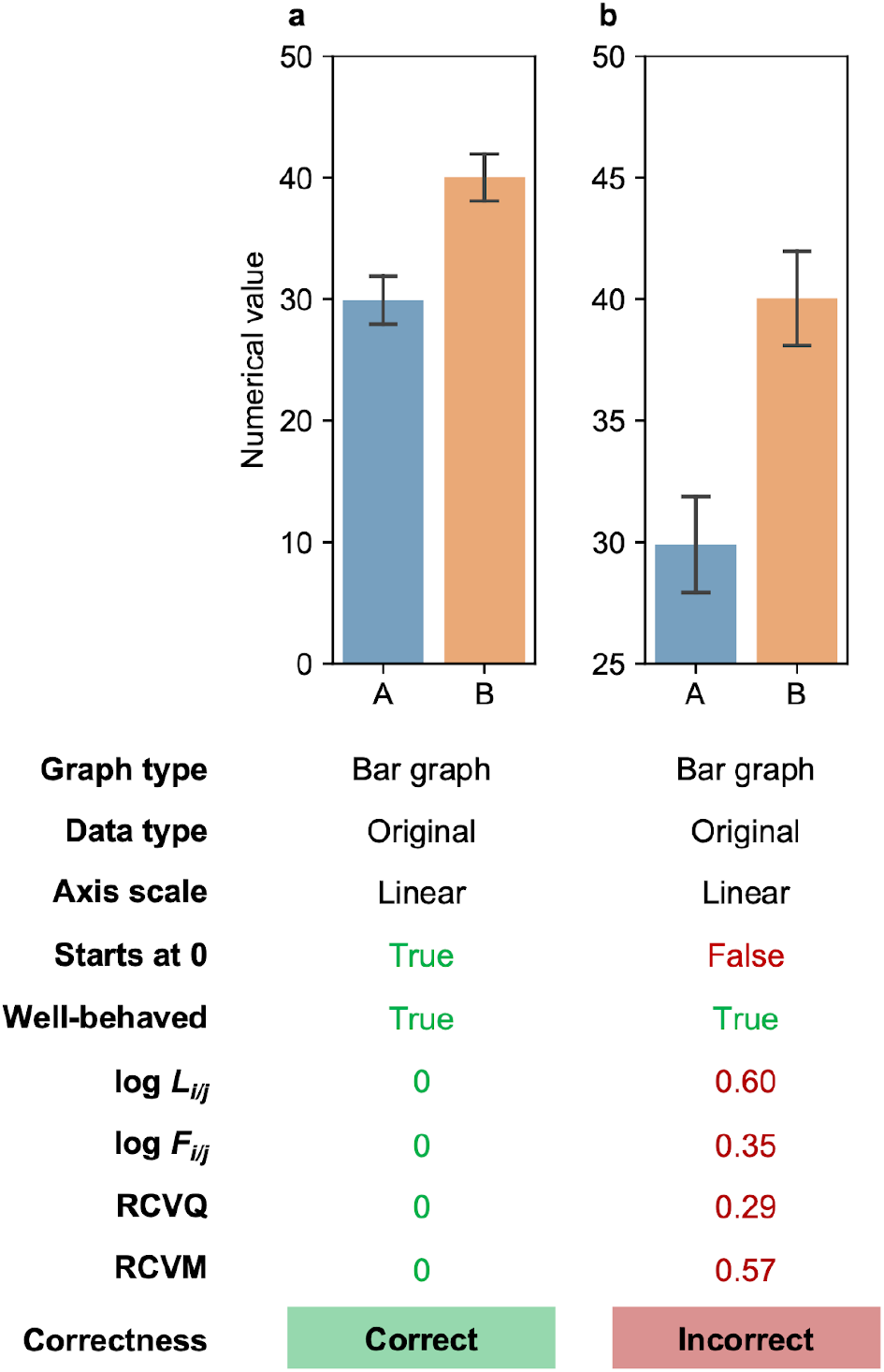
Visualizing data on a linear scale using bar graphs. All graphs visualize the same data generated from two *n* = 1000 samples from normal distributions with means *μ*_A_ = 30 and *μ*_B_ = 40 and standard deviations *σ*_A_ = *σ*_B_ = 2. Data could represent numerical values or ratios. Raw data is not shown for clarity. (**a**) Correct bar graph has y-axis starting at zero. (**b**) Incorrect bar graph has y-axis starting at nonzero values. NaN, not a number (undefined due to value not in the domain of logarithmic function).

**Extended Data Fig. 2.**
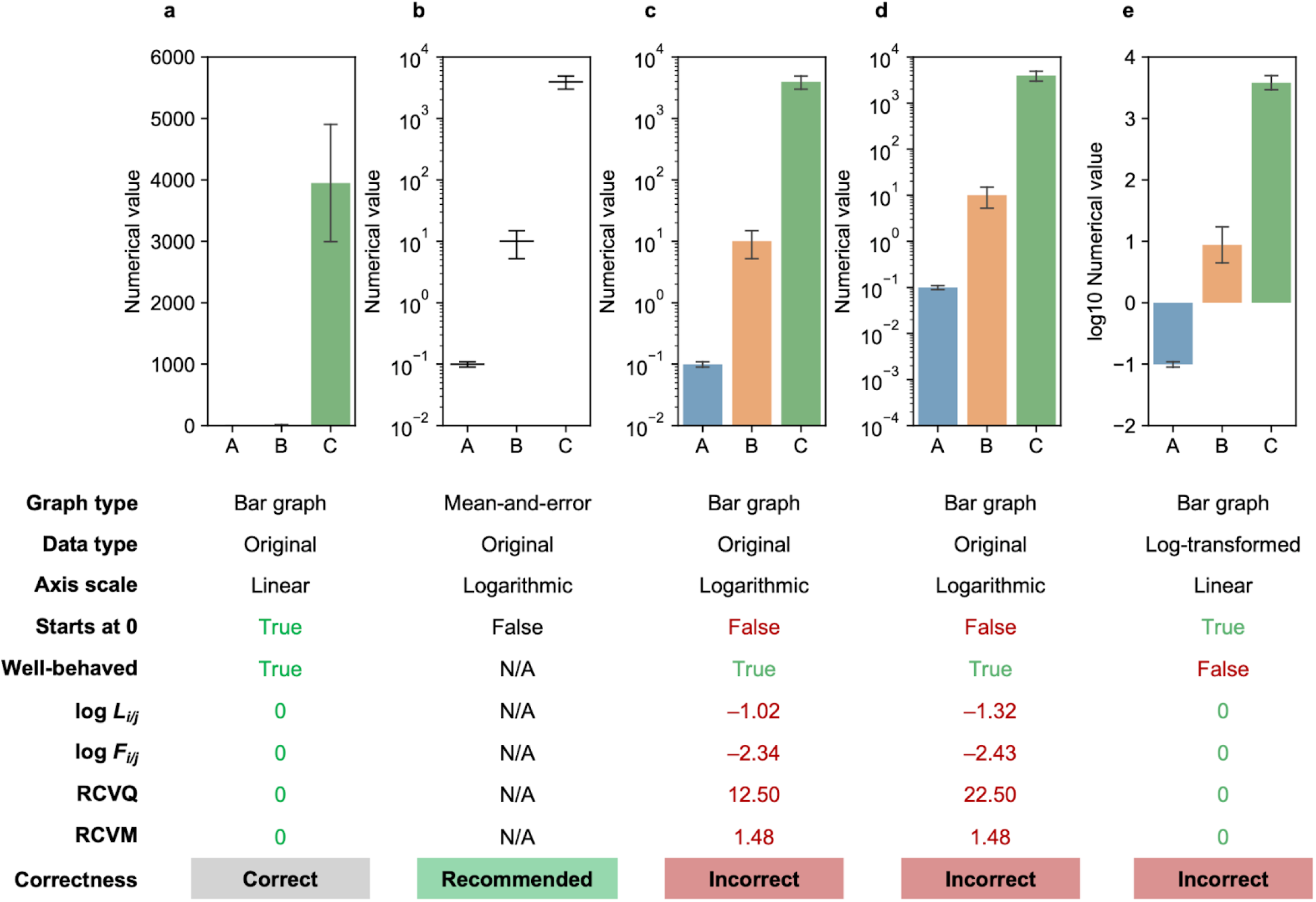
Visualizing measured values with logarithmic (order of magnitude) difference. All graphs visualize the same data generated from three *n* = 1000 samples from normal distributions with means *μ*_A_ = 0.1, *μ*_B_ = 10, and *μ*_C_ = 4000 and standard deviations *σ*_A_ = 0.01, *σ*_B_ = 5, and *σ*_C_ = 1000. Data represents numerical values, but not ratios, such as percentage, concentration, and count. Raw data is not shown for clarity. (**a**) Bar graph on a linear scale correctly visualizes the data but fails to distinguish values of different orders of magnitude. (**b**) Mean-and-error plot correctly visualizes the data and distinguishes values of different orders of magnitude. (**c, d**) Bar graphs on a logarithmic scale incorrectly visualize data because of a lack of zero baseline. (**e**) Bar graph of log-transformed data on a linear scale incorrectly visualizes measured values because the bars’ directionality is misleading and does not have practical significance. NaN, not a number (undefined due to value not in the domain of logarithmic function); N/A, not applicable.

**Extended Data Fig. 3.**
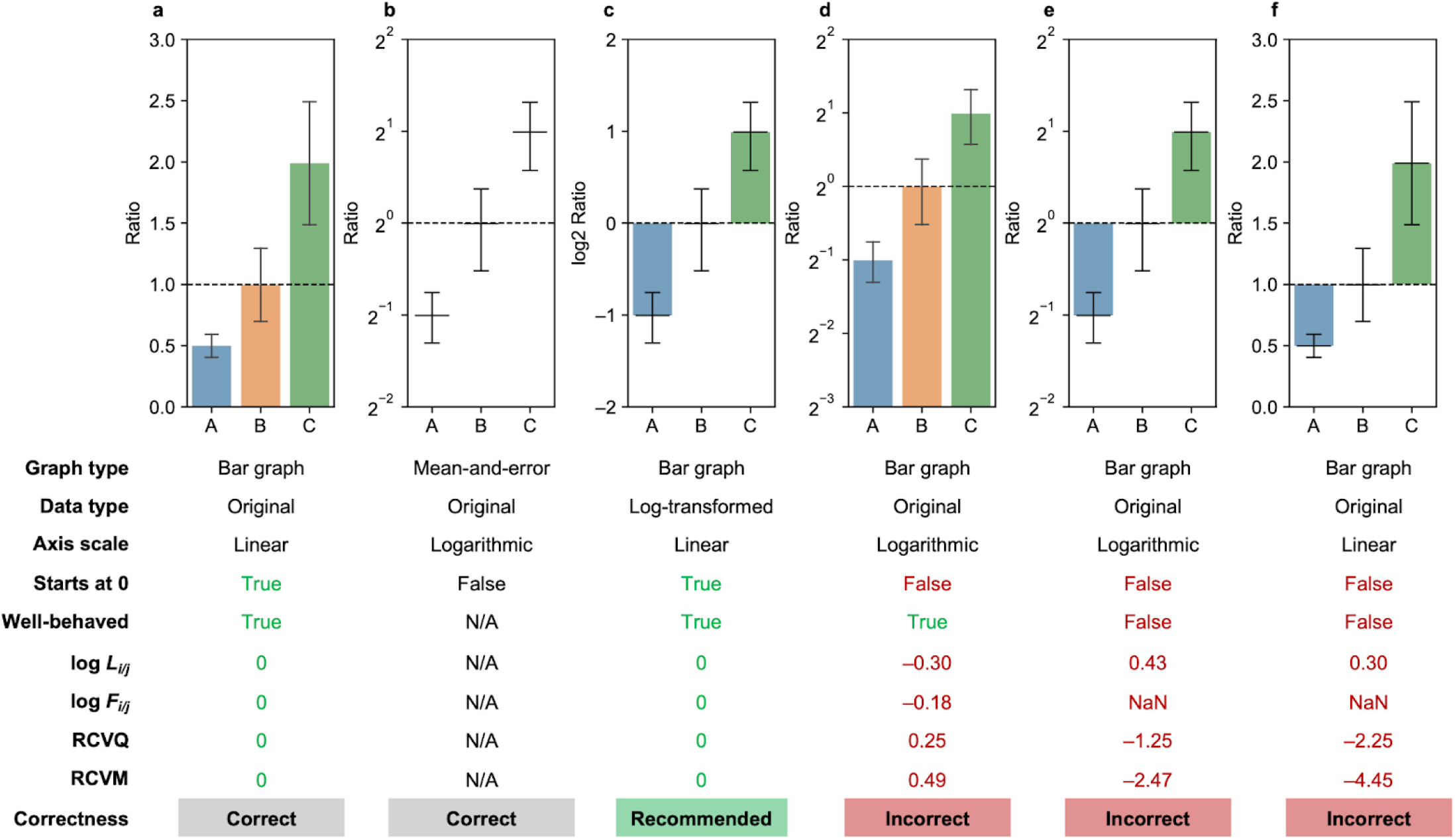
Visualizing linear fold or relative changes of ratios. All graphs visualize the same data generated from three *n* = 1000 samples from normal distributions with means *μ*_A_ = 0.5, *μ*_B_ = 1, and *μ*_C_ = 2 and standard deviations *σ*_A_ = 0.1, *σ*_B_ = 0.3, and *σ*_C_ = 0.5. Data represents ratios where the fold change between experimental groups is important, such as normalized gene expression level of wild type and partial knockout. Raw data is not shown for clarity. (**a**) Bar graph on a linear scale correctly visualizes the magnitude of absolute changes but fails to distinguish the magnitude of fold changes between ratios in different experimental groups. (**b**) Mean-and-error plot correctly visualizes the data but is hard to compare the fold changes. (**c**) Bar graph of log-transformed data on a linear scale effectively visualizes the fold changes in ratios from different experimental groups. (**d**) Bar graph on a logarithmic scale incorrectly visualizes data because of a lack of zero baseline. (**e**) Bar graph on a logarithmic scale with a baseline of 2^0^ and (**f**) bar graph on a linear scale with a baseline of 1 incorrectly visualize the data because the area of each bar does not match the measurand represented by the y-axis label (i.e. labeled ratio, but bars represent log2 ratio in **d** and (ratio – 1) in **e**). NaN, not a number (undefined due to value not in the domain of logarithmic function); N/A, not applicable; Inf, infinity.

**Extended Data Fig. 4.**
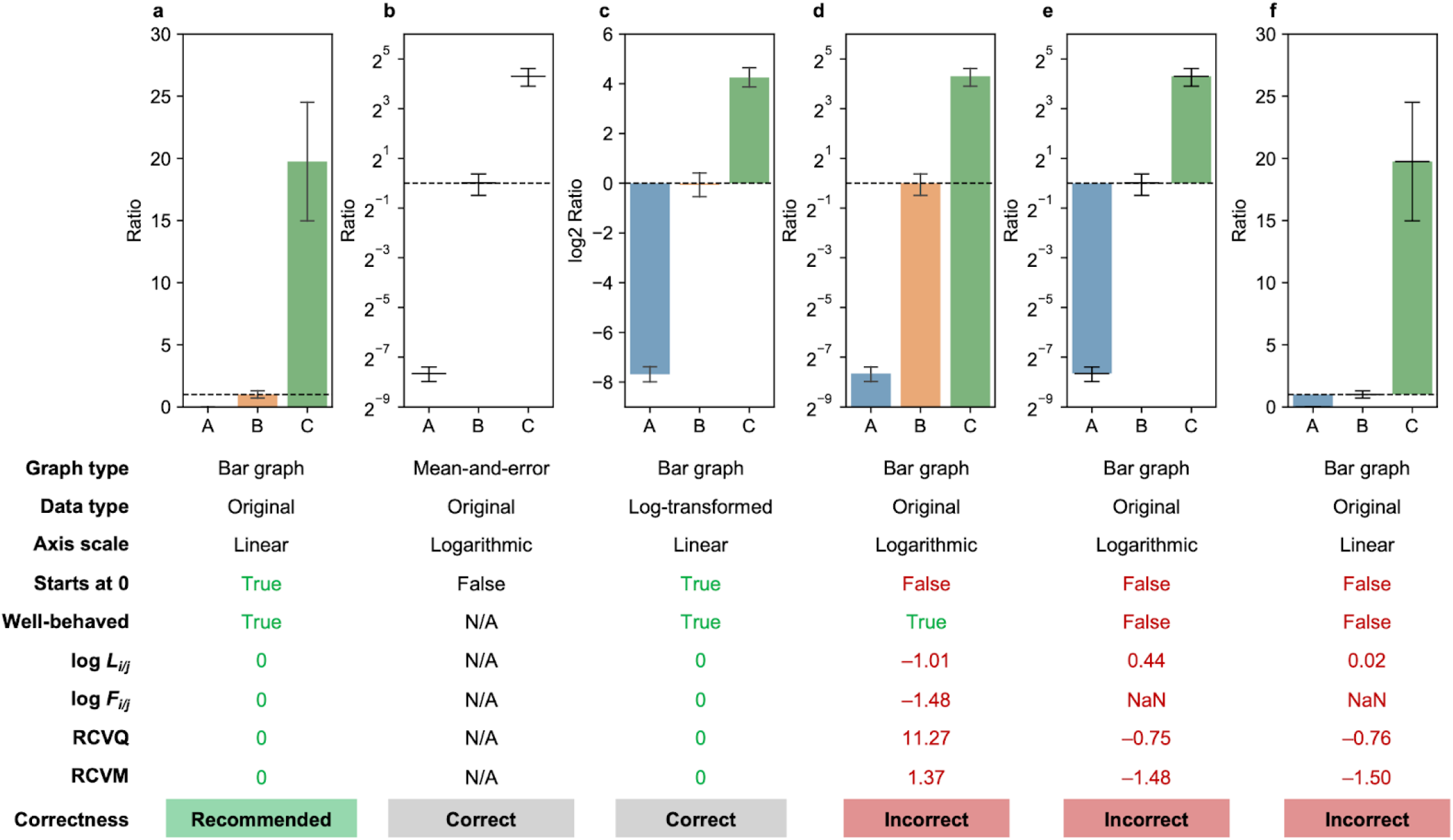
Visualizing absolute difference of ratios with near-zero values. All graphs visualize the same data generated from three *n* = 1000 samples from normal distributions with means *μ*_A_ = 0.005, *μ*_B_ = 1, and *μ*_C_ = 20 and standard deviations *σ*_A_ = 0.001, *σ*_B_ = 0.3, and *σ*_C_ = 5. Data represents ratios where absolute change and showing near-zero values is important, such as normalized gene expression levels of zero background and wild type. Raw data is not shown for clarity. (**a**) Bar graph on a linear scale effectively visualizes absolute changes in ratios and near-zero value of group A. (**b**) Mean-and-error plot on a logarithmic scale of original data and (**c**) bar graph on a linear scale of log-transformed data correctly visualize the data but are ineffective in showing absolute changes in ratios and near-zero value of group A. (**d**) Bar graph on a logarithmic scale incorrectly visualizes data because of a lack of zero baseline. (**e**) Bar graph on a logarithmic scale with a baseline of 2^0^ and (**f**) bar graph on a linear scale with a baseline of 1 incorrectly visualize the data because the area of each bar does not match the measurand represented by the y-axis label (i.e. labeled ratio, but bars represent log2 ratio in **d** and (ratio - 1) in **e**). NaN, not a number (undefined due to value not in the domain of logarithmic function); N/A, not applicable; Inf, infinity.

**Extended Data Fig. 5.**
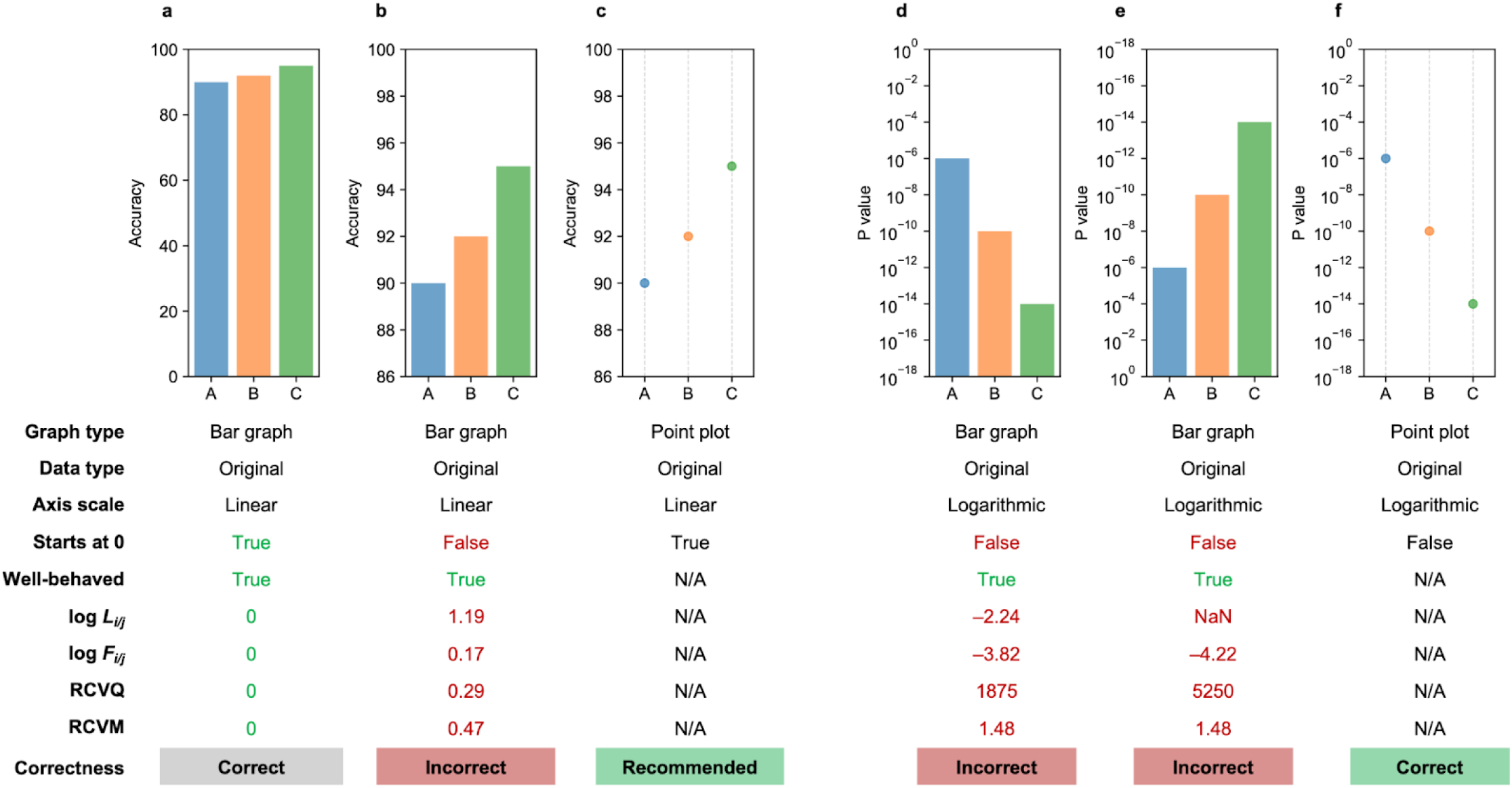
Visualizing single sample values in linear and log scales. (**a**-**c**) All graphs visualize the same simulated data where absolute changes are important rather than relative change or fold change, such as prediction metrics, on a linear scale. (**a**) Bar graph on a linear scale starting at zero correctly visualizes the data but does not emphasize the absolute change. (**b**) Bar graph not starting at zero distorts the data. (**c**) Point plot on a linear scale correctly visualizes the data and emphasizes the absolute change. (**d**-**f**) All graphs visualize the same simulated *P* value data plotted on a logarithmic scale. *P* values are usually incorrectly plotted with (**d**) bar graph on a logarithmic scale starting at an arbitrary value, or (**e**) bar graph starting at 10^0^, where the bars attempt to represent the relative level of significance but not *P* value. Note in **e**, *P* value nonintuitively decreases from bottom to top. (**f**) Point plot on a logarithmic scale correctly visualizes *P* values. NaN, not a number (undefined due to value not in the domain of logarithmic function); N/A, not applicable.

**Extended Data Fig. 6.**
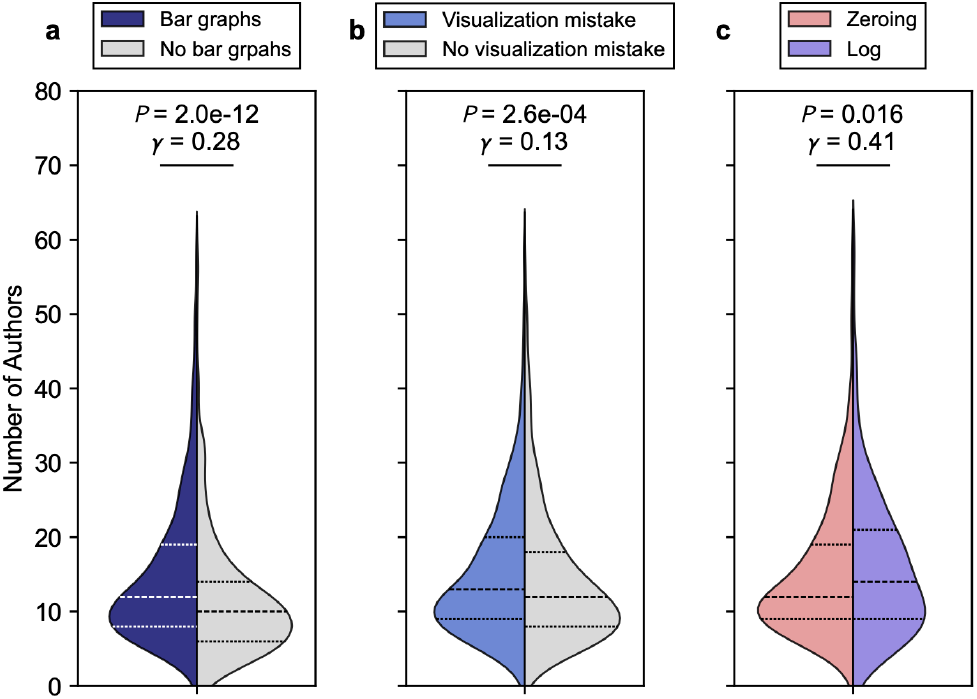
Articles with bar graph visualization mistakes has more authors. Comparison of author number distribution of (**a**) articles with (*n* = 2985) and without (*n* = 402) bar graphs, (**b**) articles with bar graphs that have (*n* = 873) and do not have (*n* = 2112) incorrect bar graph visualization, and (**c**) articles with zeroing (*n* = 524) and log (*n* = 284) mistakes. Articles with >60 authors (<1.5% of the population) are not plotted for clarity. Dashed lines represent 25, 50, and 75 percentiles. *P* values were computed using two-sided Mann-Whitney U test. Effect sizes were computed using nonparametric gamma.

**Extended Data Fig. 7.**
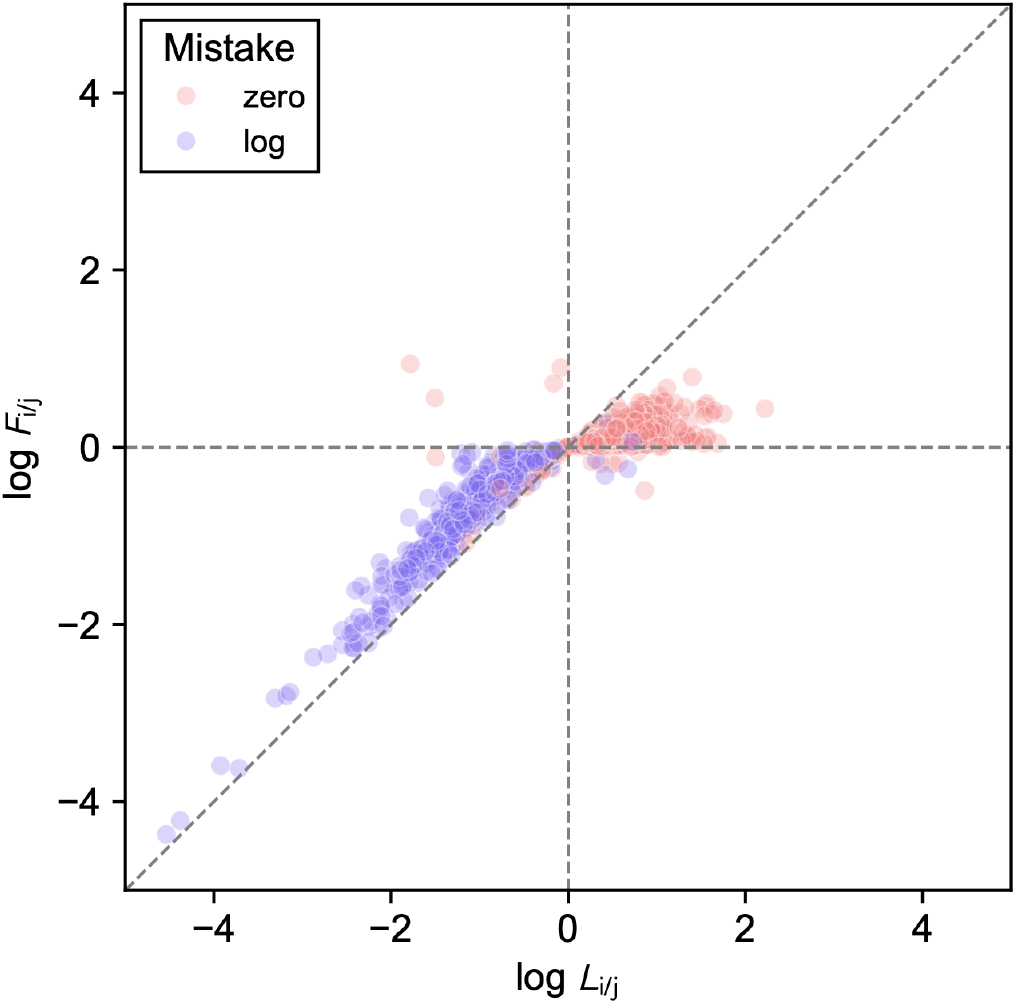
Correlation between lie factor of relative change and fold change. Scatter plot of bias-mitigated log-transformed lie factor of fold change *F*_*i/j*_ with respect to that of relative change *L*_*i/j*_ hued with zeroing (*n* = 722) and log (*n* = 392) mistakes. Diagonal dashed line represents the identity function (*y* = *x*).

**Extended Data Fig. 8.**
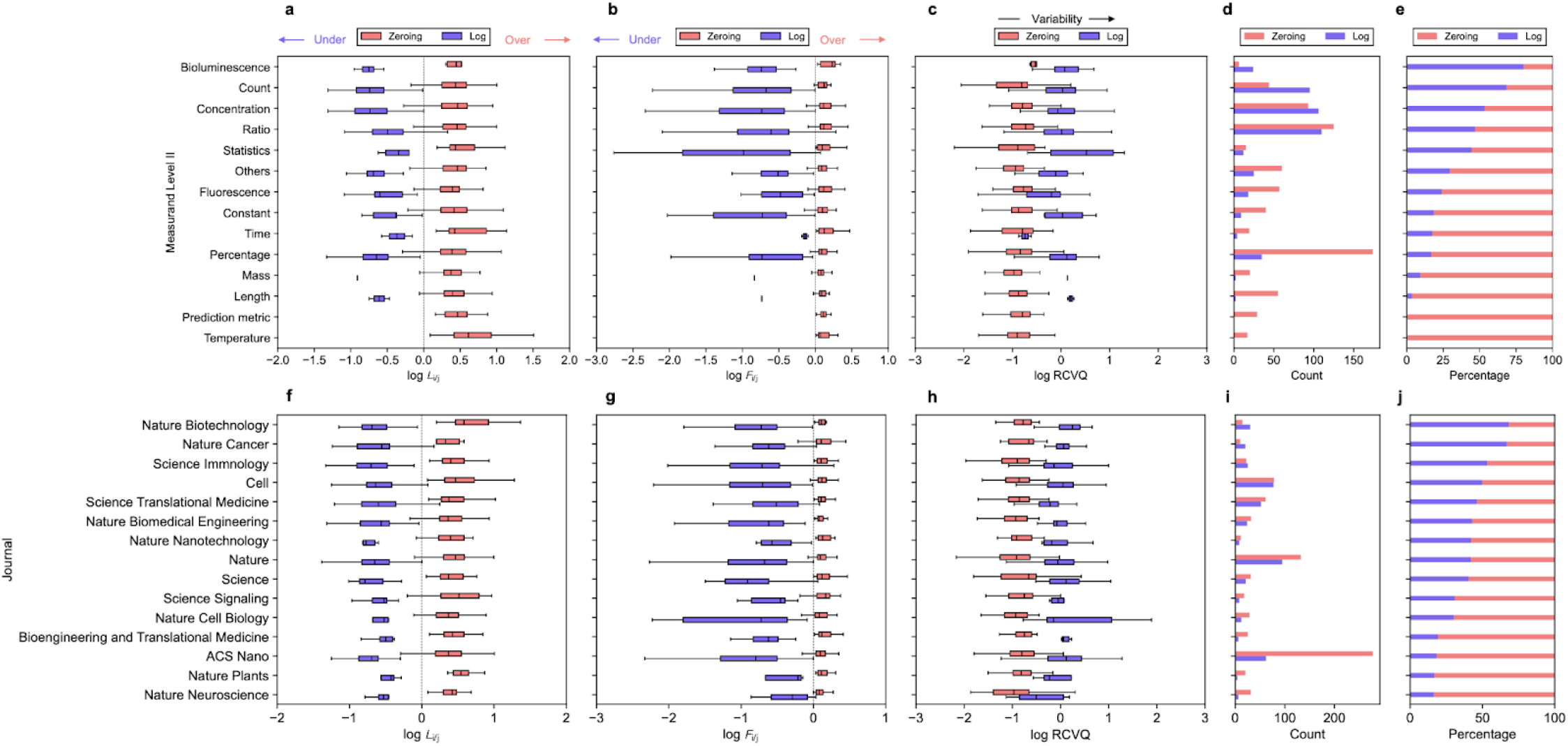
Extent and prevalence of data distortion in bar graphs by measurand represented and by journal. Bar graphs with zeroing and log mistakes are grouped by measurand represented (**a**-**e**) or journal (**f**-**j**). Box plot of bias-mitigated log-transformed (**a, f**) lie factor of relative change *L*_*i/j*_, (**b, g**) lie factor of fold change *F*_*i/j*_, and (**c, h**) RCVQ. (**d, i**) Count and (**e, j**) percentage of zeroing and log mistakes. Box plot shows 25, 50, and 75 percentiles. Whiskers extend to the farthest data point within 1.5 times the interquartile range.

**Extended Data Fig. 9.**
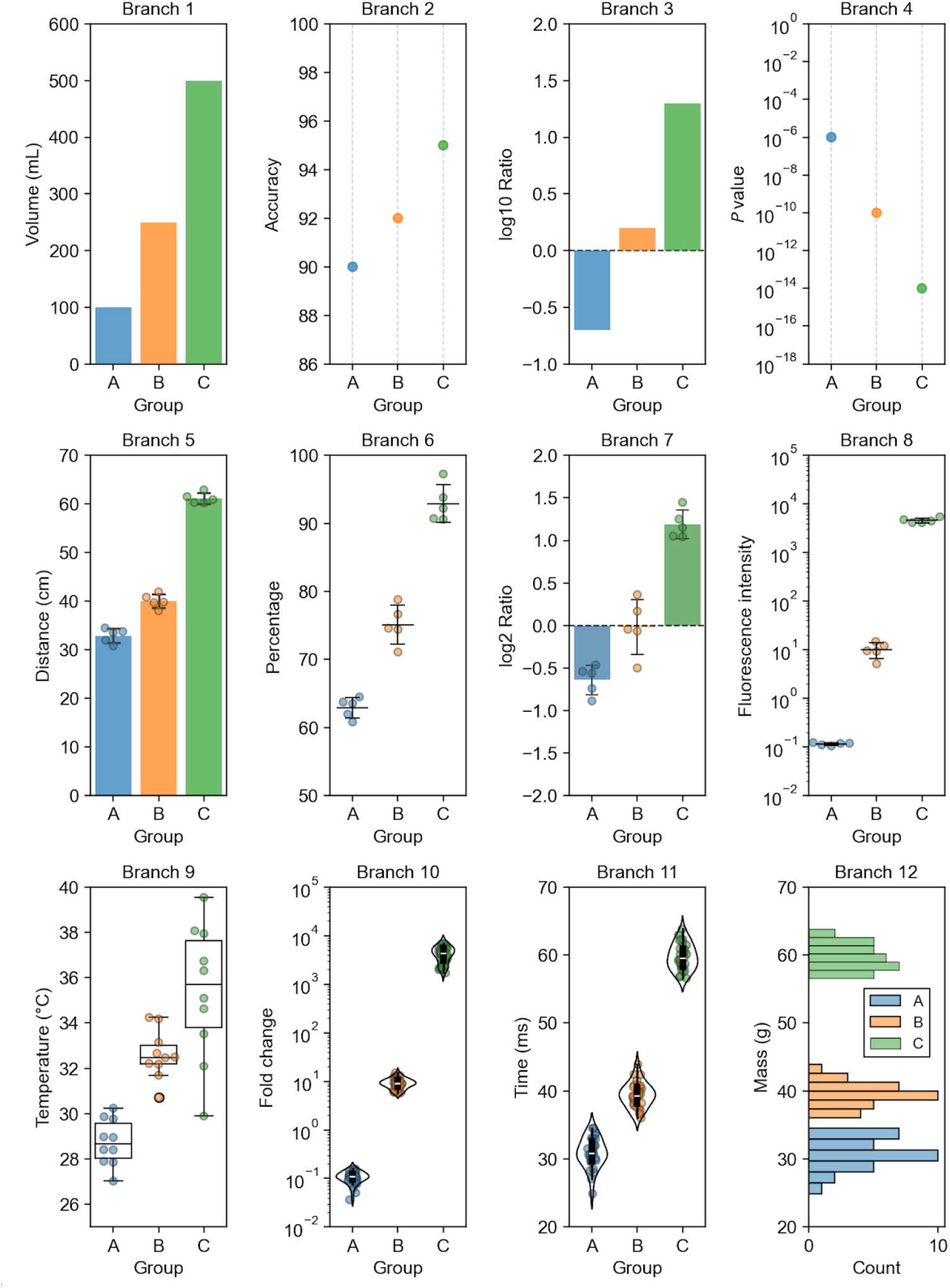
Examples of graphs for each branch in the decision tree in Fig. 2d with simulated data.

**Extended Data Fig. 10.**
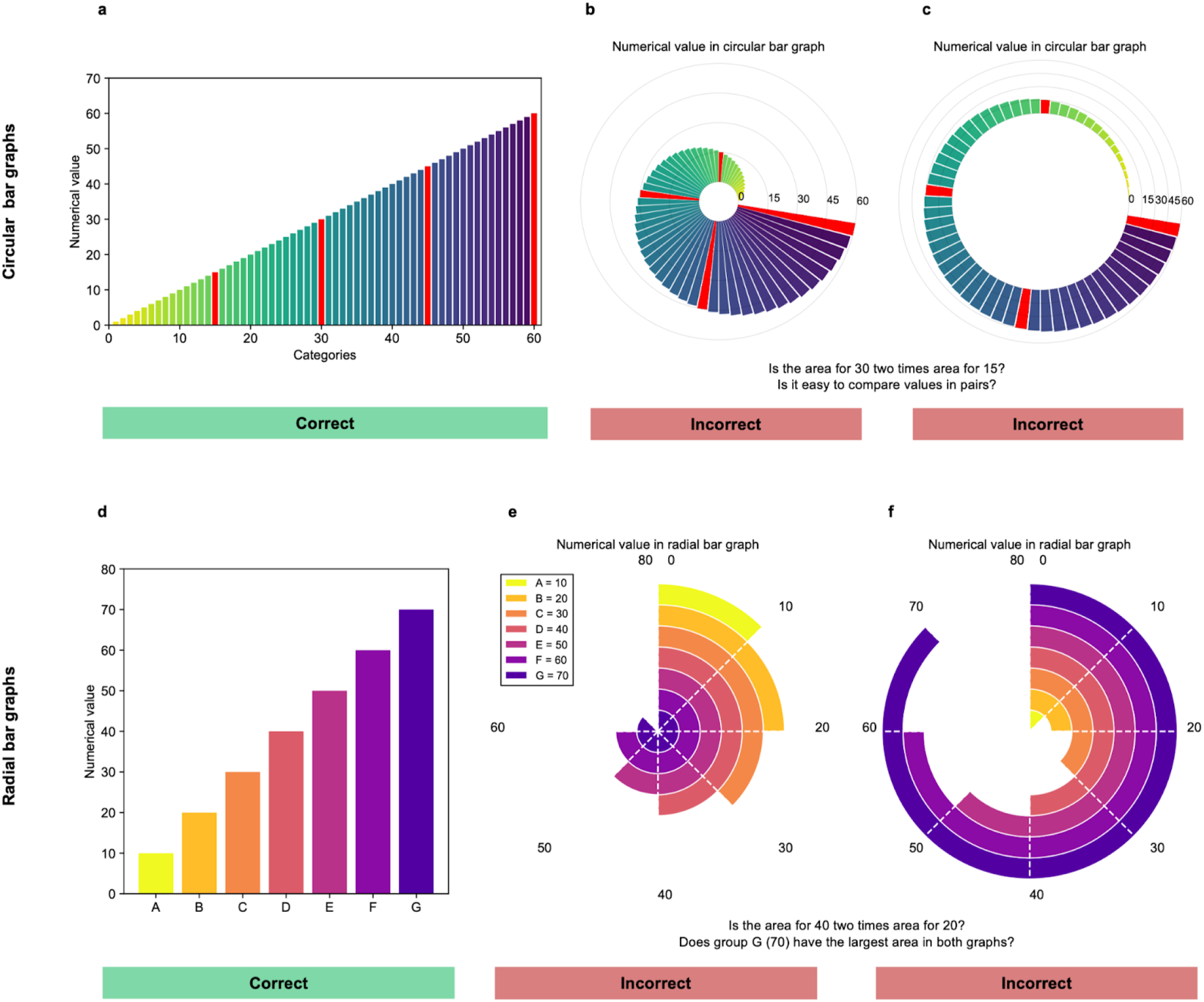
Bar graphs in polar coordinates inherently distort the underlying data and should not be used. (**a**-**c**) The same simulated data from 0 to 60 with an increment of 1 demonstrates the distortion caused by circular bar graphs. Circular bar graphs use the angular coordinate to represent nominal variables and the radial coordinate to represent numerical variables. Red bars represent values of 15, 30, 45, and 60. (**a**) Bar graph in Cartesian coordinates correctly represents the data. (**b, c**) Circular bar graphs distort the data to different extents depending on the inner radius at which the bar starts. The lack of alignment between bars also makes it hard to make pairwise comparisons. (**d**-**f**) The same simulated data from 10 to 70 with an increment of 10 demonstrates the distortion caused by radial bar graphs. Radial bar graphs use the radial coordinate to represent nominal variables and the angular coordinate to represent numerical variables. (**d**) Bar graph in Cartesian coordinates correctly represents the data. (**e, f**) Radial bar graphs arbitrarily distort the data based on the radial coordinate at which the experimental group is placed.

