## Supplementary Information for "Quantifying Data Distortion in Bar Graphs in Biological Research"

### Supplementary Information
### Quantifying Data Distortion in Bar Graphs in Biological Research

Teng-Jui Lin, Markita P. Landry

October 4, 2024

### 1 Supplementary Methods

#### 1.1 Search Queries

Search queries from the publisher's website used to acquire the article's PDF are listed below. To quantify only research articles in biological sciences and engineering, multidisciplinary journals are subjected to additional filtering.

*Nature Nanotechnology* search is constrained by the subject of "Biotechnology."

*Nature* search is individually constrained by the subjects of "Cancer," "Cell Biology," "Molecular Biology," "Biotechnology," "Neuroscience," "Drug Discovery," "Chemical Biology," "Microbiology," "Immunology," "Stem Cells," "Genetics," "Computational Biology and Bioinformatics," "Developmental Biology," "Diseases," "Evolution," "Medical Research," and "Plant Sciences." Articles present in at least one of the subjects (OR filtering) are included.

*Science* search is constrained by search terms of "Cancer," "Cell Biology," "Molecular Biology," "Biotechnology," "Neuroscience," "Drug Discovery," "Chemical Biology," "Microbiology," "Immunology," "Stem Cells," "Genetics," "Computational Biology and Bioinformatics," "Developmental Biology," "Diseases," "Evolution," "Medical Research," and "Plant Sciences" using OR filtering. Irrelevant articles are excluded manually.

*ACS Nano* search is constrained by the topic of "Biology and Biological Chemistry."

The links to search queries are provided below.

- *Science Translational Medicine*
- *Science Immunology*
- *Science Signaling*
- *Nature Biotechnology*
- *Nature Nanotechnology* (by subject)
- *Nature Biomedical Engineering*
- *Nature Cancer*
- *Nature Plants*
- *Nature Cell Biology*
- *Nature Neuroscience*
- *Bioengineering & Translational Medicine*
- *Cell*
- *Nature* (by subject)
  - Cancer
  - Cell Biology
  - Molecular Biology
  - Biotechnology
  - Neuroscience
  - Drug Discovery
  - Chemical Biology
  - Microbiology
  - Immunology
  - Stem cells

- Genetics
  - Computational biology and bioinformatics
  - Developmental biology
  - Diseases
  - Evolution
  - Medical Research
  - Plant Sciences

- *Science* (By search term)
- *ACS Nano*

##### 1.2 Measurand annotations

Four categorizations of measurand annotations at different scopes provide different levels of granularity of the quantities being represented in bar graphs. The elements within a categorization are mutually exclusive. The following list presents the categorizations. The numbers in the parentheses are the number of elements within each categorization scheme.

- Absolute/Relative (2): “Absolute”, “Relative”
- Measurement Type (4): “Measured value”, “Ratio”, “Percentage”, “Calculated value”
- Measurand Level II (14): “Temperature”, “Statistics”, “Mass”, “Time”, “Bioluminescence”, “Constant”, “Length”, “Others”, “Fluorescence”, “Prediction metric”, “Count”, “Percentage”, “Ratio”, “Concentration”
- Measurand Level I (34): “Total radiant efficiency”, “Radiance”, “Energy”, “Radiant efficiency”, “Angle”, “Statistics”, “Potential”, “Particle diameter”, “Area”, “Radiant flux”, “Volume”, “Colon length”, “Cell viability”, “p-value”, “Index”, “Temperature”, “Constant”, “Prediction metric”, “Mass”, “Length”, “Time”, “Accuracy”, “Characteristic concentration”, “Others”, “AUC”, “Cytokine concentration”, “Cell percentage”, “Concentration”, “Cell count”, “Relative luminescence”, “Count”, “Percentage”, “Titer”, “Ratio”

##### 1.3 Statistics

A nonparametric Cohen’s d-consistent effect size $\gamma$ [1] is used to estimate effect size, where we’re interested in the median (0.5 quantile, $p = 0.5$):

$$\gamma \equiv \gamma_p = \frac{Q_p(Y) - Q_p(X)}{\mathrm{PMAD}_{XY}},$$

where $Q_p$ is the $p^{\text{th}}$ quantile of a given sample, $X$ and $Y$ the two samples of interest, and $\mathrm{PMAD}_{XY}$ is the pooled median absolute deviation defined as

$$\mathrm{PMAD}_{XY} = \sqrt{\frac{(n_X - 1)\mathrm{MAD}_X^2 + (n_Y - 1)\mathrm{MAD}_Y^2}{n_X + n_Y - 2}},$$

where $n_X$ and $n_Y$ are the sample size of $X$ and $Y$, and MAD is the median absolute deviation defined as

$$\mathrm{MAD}_X = C\ \mathrm{median}(|X_i - \mathrm{median}(X)|),$$

where $C = 1.4826$ is the consistent constant.

### 2 Supplementary Discussion

#### 2.1 Fundamentals of Data Visualizations

##### 2.1.1 Principle of proportional ink

To represent data value in a graphical form, we need to create graphical elements called **marks** [2], such as points, lines, and areas. The appearance of these marks, or **channels**, can be used to encode information about the data, whether it's about its identity (categorical) or magnitude (numerical) [2]. For example, a bar graph uses bars as marks and uses the bars' area as a magnitude channel to encode the data value.

In 1983, Edward Tufte proposed the **principle of proportional ink**, stating that "the representation of numbers, as physically measured on the surface of the graphic itself, should be directly proportional to the numerical quantities represented" [3]. In other words, the data value (**true value**) must be directly proportional to the magnitude channel (**visualized value**) of their respective marks for a **well-represented** graph. For each pair of true and visualized values, the proportionality constants should all be the same.

##### 2.1.2 Mathematical definitions of data visualizations

**2.1.2.1 Mark proportionality constants** Suppose we have a **true value vector** $\mathbf{x} = [x_1, x_2, ..., x_n]^T$ that consists of $n$ marks' true values of $x_i \in \mathbb{R}$, which we visualize using a mark's magnitude channel with visualized values of $y_i \in \mathbb{R}$ that are elements of the **visualized value vector** $\mathbf{y} = [y_1, y_2, ..., y_n]^T$. Each mark has a **mark proportionality constant** between the true value and the visualized value

$$\alpha_i \equiv y_i/x_i, \tag{2.1}$$

which are elements of the **mark proportionality constant vector** $\boldsymbol{\alpha}$ that can be defined by an element-wise division $\oslash$:

$$\boldsymbol{\alpha} = \mathbf{y} \oslash \mathbf{x}. \tag{2.2}$$

The proportionality relation between visualized value and true value can therefore be expressed as an element-wise multiplication $\odot$:

$$\mathbf{y} = \boldsymbol{\alpha} \odot \mathbf{x}. \tag{2.3}$$

Mathematically, the principle of proportionality suggests that a well-represented graph has all its mark proportionality constants the same: $\alpha_1 = \alpha_2 = \cdots = \alpha_n \equiv \alpha$, so the mark proportionality constant reduces to

$$\mathbf{y} = \alpha\mathbf{x}, \tag{2.4}$$

where $\alpha$ is the **graph proportionality constant** of a well-represented graph.

**2.1.2.2 Theoretical visualized values** For each mark, we can define theoretical visualized values if they were to be well-represented based on some mark. Mathematically, the **theoretical visualized value** for mark $i$ based on the mark proportionality constant of mark $j$ is defined by

$$y'_{i/j} \equiv x_i\alpha_j. \tag{2.5}$$

The **theoretical visualized value vector** of all marks in a graph based on some mark $j$ is

$$\mathbf{y}'_{/j} = \alpha_j\mathbf{x}. \tag{2.6}$$

Similarly, the theoretical visualized value vector of mark $i$ in a graph based on all marks is

$$\mathbf{y}'_{i/} = x_i\boldsymbol{\alpha}. \tag{2.7}$$

If we compute all the theoretical visualized values based on all $n$ marks, we can construct the **theoretical visualized value matrix**

$$\mathbf{Y}' = \begin{bmatrix} \mathbf{y}'_{/1} & \mathbf{y}'_{/2} & \cdots & \mathbf{y}'_{/n} \end{bmatrix} \tag{2.8}$$

$$= \begin{bmatrix} \mathbf{y}'_{1/} & \mathbf{y}'_{2/} & \cdots & \mathbf{y}'_{n/} \end{bmatrix}^T \tag{2.9}$$

$$= \begin{bmatrix} y'_{1/1} & y'_{1/2} & \cdots & y'_{1/n} \\ y'_{2/1} & y'_{2/2} & \cdots & y'_{2/n} \\ \vdots & \vdots & \ddots & \vdots \\ y'_{n/1} & y'_{n/2} & \cdots & y'_{n/n} \end{bmatrix}, \tag{2.10}$$

where column $i$ represents the theoretical visualized values of each mark based on mark $i$, and row $j$ represents the theoretical visualized values of mark $j$ based on each mark. The matrix can be obtained with

$$\mathbf{Y}' = \boldsymbol{\alpha}\mathbf{x}^T. \tag{2.11}$$

**2.1.2.3 Difference proportionality constants** We can similarly define proportionality constants for value differences. For a true value difference $x_i - x_j$ that is represented with visualized value difference $y_i - y_j$, the **difference proportionality constant** is defined as

$$\alpha_{i/j} = \frac{y_i - y_j}{x_i - x_j}. \tag{2.12}$$

Note that the difference proportionality constant is defined on a pairwise combination level because it is order invariant:

$$\alpha_{i/j} = \alpha_{j/i}. \tag{2.13}$$

For a well-represented graph, the difference proportionality constant is the same as the mark proportionality constants and graph proportionality constant: $\alpha_{i/j} = \alpha_i = \alpha$.

If the visualized value is linearly proportional to the true value ($y(x) = kx + b$), we have

$$\alpha_{i/j} = \frac{y_i - y_j}{x_i - x_j} = \frac{(kx_i + b) - (kx_j + b)}{x_i - x_j} = k, \tag{2.14}$$

suggesting that the difference proportionality constant is a constant regardless of which value difference is selected for calculation. The difference proportionality constant reduces to a graph-level property.

Interestingly, the difference proportionality constant is the same for linearly proportional ($b \neq 0$, misrepresented) and directly proportional ($b = 0$, well-represented) relationships. However, only directly proportional relationships satisfy the principle of proportional ink, suggesting that the difference proportionality constant for a linearly proportional relationship can serve as the ground truth graph proportionality constant if each mark were to be well-represented:

$$\alpha_{i/j} = k \equiv \alpha. \tag{2.15}$$

If the visualized value is linearly proportional to the logarithm of the true value ($y(x) = k\log(x) + b$), we have

$$\alpha_{i/j} = \frac{y_i - y_j}{x_i - x_j} = \frac{(k\log(x_i) + b) - (k\log(x_j) + b)}{x_i - x_j} = \frac{k}{x_i - x_j}\log\left(\frac{x_i}{x_j}\right), \tag{2.16}$$

which is not a constant and depends on the value pair selected.

#### 2.2 Metrics to Quantify Data Distortion

When quantifying the extent to which data visualization distorts the underlying data, we need to recognize that the distortion is based on *comparisons* between marks rather than stand-alone visualizations with one mark. Any stand-alone visualization with one mark can have an arbitrary visualized value based on the principle of proportional ink. The scaling of such stand-alone visualization does not affect how we interpret the data. For example, for a bar graph with only one bar, the length of the bars does not matter for interpretation. However, when comparing between two marks, the principle of proportional ink demands that the mark and proportionality constants be the same. Because comparison between marks is meaningful but not the visualized value of a mark, we only consider graphs with $\geq 2$ marks to quantify data distortion (i.e. $\mathbf{x}$ and $\mathbf{y}$ must have at least 2 elements). We also only consider non-zero visualized values ($y_i \neq 0$) because marks with zero visualized values cannot be meaningfully compared with other marks.

In the following sections, we establish a rigorous mathematical framework for data distortion metrics that are previously developed (but not rigorously characterized) or developed in this work.

##### 2.2.1 Tufte's Lie Factor

**2.2.1.1 Mathematical definition of lie factors** To quantify the extent to which the graphs' data are distorted, Edward Tufte in 1983 defined the **lie factor (of relative change)** [3] as

$$\text{Lie factor} = \frac{\text{relative change of visualized value}}{\text{relative change of true value}}, \tag{2.17}$$

where the relative change is:

$$\text{Relative change} = \frac{\text{Value 2} - \text{Value 1}}{\text{Value 1}}. \tag{2.18}$$

Some people define relative change with absolute values. However, we do not adopt such convention because the negative signs have practical significance, such as encoding for the directionality of bar graphs or negative true values.

Because relative change is normalized based on one of the values, the lie factor is also normalized with the same basis. We can therefore define two lie factors for each pairwise relative comparison. Mathematically, we can define the lie factor of the relative change between mark $i$ and mark $j$'s magnitude channel normalized to mark $j$'s magnitude channel ($i \neq j$) as

$$L_{i/j} \equiv \frac{E_{yi/j}}{E_{xi/j}}, \tag{2.19}$$

where we define the size of effect of the visualized and true value as

$$E_{yi/j} \equiv \frac{y_i - y_j}{y_j} \quad \text{and} \quad E_{xi/j} \equiv \frac{x_i - x_j}{x_j}, \tag{2.20}$$

respectively. Expand and simplify the lie factor, we have

$$L_{i/j} = \frac{x_j}{y_j}\frac{y_i - y_j}{x_i - x_j} = \frac{\alpha_{i/j}}{\alpha_j}, \tag{2.21}$$

suggesting that the lie factor is the ratio between the difference proportionality constant between marks $i$ and $j$ and the mark proportionality constant of the normalized mark. We can similarly define the lie factor of the relative change between mark $i$ and mark $j$'s magnitude channel normalized to mark $i$'s magnitude channel as

$$L_{j/i} \equiv \frac{E_{yj/i}}{E_{xj/i}} = \frac{\alpha_{j/i}}{\alpha_i}, \tag{2.22}$$

with the size of effect also similarly defined. Note that the pairwise lie factors normalized to different marks are related by the mark proportionality constants by

$$\frac{L_{i/j}}{L_{j/i}} = \frac{\alpha_i}{\alpha_j}, \tag{2.23}$$

if the true value difference and visualized value difference are nonzero ($x_i \neq x_j$ and $y_i \neq y_j$).

**2.2.1.2 Behavior of the lie factor** If the graph's marks are well-represented according to the principle of proportional ink [3] such that their mark proportionality constants are the same as the difference proportionality constants, the lie factor simplifies to unity:

$$L_{i/j} = \frac{\alpha_{i/j}}{\alpha_j} = 1, \tag{2.24}$$

$$L_{j/i} = \frac{\alpha_{j/i}}{\alpha_i} = 1. \tag{2.25}$$

What happens when the graph is misrepresented? If the visualized values over-represent the true value, the lie factors are greater than 1; if the visualized values under-represent the true value, the lie factors are positive and less than 1. Let's consider a representative example for analysis.

For the following analysis, consider misrepresented but "well-behaved" graphs that (1) have positive values ($x_i > 0, x_j > 0$) and (2) do not visualize positive values with negative magnitude channels and vice versa ($\text{sgn}(x_i) = \text{sgn}(y_i)$, $\text{sgn}(x_j) = \text{sgn}(y_j)$). Consider the case of $x_i > x_j > 0$, $y_i > y_j > 0$. To over-represent the true value, either $y_j$ decreases by $k_j > 0$ or $y_i$ increases by $k_i > 0$ in the over-represented lie factor $L'_{i/j}$:

$$\begin{aligned} L'_{i/j} &= \frac{y'_i \uparrow - y'_j \downarrow}{y'_j \downarrow} \frac{x_j}{x_i - x_j} \\ &= \frac{(y_i + k_i) - (y_j - k_j)}{y_j - k_j} \frac{x_j}{x_i - x_j} \end{aligned}$$

We ask if the over-represented lie factor is greater than the well-represented lie factor $L_{i/j} = 1$:

$$\begin{aligned} L'_{i/j} &\overset{?}{>} L_{i/j} \\ \frac{(y_i + k_i) - (y_j - k_j)}{y_j - k_j} \frac{x_j}{x_i - x_j} &\overset{?}{>} \frac{y_i - y_j}{y_j} \frac{x_j}{x_i - x_j} \\ \frac{y_i - y_j + k_i + k_j}{y_j - k_j} &\overset{?}{>} \frac{y_i - y_j}{y_j} \\ y_i y_j - y_j^2 + k_i y_j + k_j y_j &\overset{?}{>} y_i y_j - y_j^2 - y_i k_j + y_j k_j \\ k_i y_j &> -y_i k_j \end{aligned} \qquad \blacksquare$$

Because all the quantities are positive, we have proved that $L'_{i/j} > 1$ subjected to the above assumptions. Similar logic applies for "well-behaved" graphs with negative values ($x_i < 0, x_j < 0$). Full proof for all cases is provided in Supplementary Proofs.

**2.2.1.3 Interpretation of the lie factor** In summary, for "well-behaved" graphs, we have the behavior of generic lie factor regardless of the normalization baseline $L$:

$$\begin{cases} L \in (1, \infty) & \text{over-represented} \\ L = 1 & \text{well-represented} \\ L \in (0, 1) & \text{under-represented} \end{cases} \tag{2.26}$$

Notice how the over-representation and under-representation of the graphs have different range scales. To more accurately reflect the extent of over-representation and under-representation, we should use the log-transformed lie factor so they span the same range:

$$\begin{cases} \log(L) \in (0, \infty) & \text{over-represented} \\ \log(L) = 0 & \text{well-represented} \\ \log(L) \in (-\infty, 0) & \text{under-represented} \end{cases} \tag{2.27}$$

By rearranging the lie factor equation, we have three interpretations for the lie factor. The lie factor quantifies the ratio between relative change in visual value and relative change in true value:

$$L_{i/j} \equiv \frac{E_{yi/j}}{E_{xi/j}}. \tag{2.19}$$

The lie factor also quantifies the ratio between the difference proportionality constant between marks $i$ and $j$ and the mark proportionality constant of the normalized mark:

$$L_{i/j} = \frac{\alpha_{i/j}}{\alpha_j}. \tag{2.21}$$

For graphs whose visual and true values are linearly proportional, because $\alpha_{i/j}$ reduces to the graph proportionality constant $\alpha$, the lie factor quantifies the ratio between the graph proportionality constant and the mark proportionality constant (i.e. deviation from well-representation):

$$L_{i/j} = \frac{\alpha}{\alpha_j} \equiv L_j, \tag{2.28}$$

which is only dependent on mark $j$.

Note that because the lie factor is based on relative change, it does not reflect the absolute magnitude of the change. Two misrepresented graphs with large differences in the absolute magnitude of the change (visually different) may have the same lie factor representing the relative change. This requires caution when interpreting lie factors.

The preceding analysis covers "well-behaved" graphs that we encounter most of the time. However, it breaks down when graphs are misrepresented and not "well-behaved." In the following sections, consider misrepresented and not "well-behaved" graphs that may give rise to negative, zero, or undefined lie factors. Some of these anomalous behaviors are previously explored in the context of graph discrepancy index (GDI, Section 2.2.2) in a qualitative manner [4], but they are more useful to discuss in the context of lie factors with rigorous quantification.

**2.2.1.4 Negative lie factors** Negative lie factors are common for bar graphs whose direction carries information about the value's sign. Consider a misrepresented bar graph with one negative value ($x_j < 0$) and one positive value ($x_i > 0$), but it is both visually represented with positive values ($y_i > y_j > 0$). This is common when graphs with negative values are not correctly baseline at zero, but baselined at some arbitrary negative value. Consider the two lie factors normalized to different marks, and note the positive and negative quantities in the equation:

$$L_{i/j} = \frac{y_i - y_j}{y_j} \frac{x_j}{x_i - x_j} < 0, \tag{2.29}$$

$$L_{j/i} = \frac{y_i - y_j}{y_i} \frac{x_i}{x_i - x_j} > 0, \tag{2.30}$$

we get a positive and a negative lie factor normalized on different marks.

As another example, consider a misrepresented bar graph on a logarithmic scale with two positive values ($x_i > x_j > 0$), but it is baselined such that one is visualized with a negative value ($y_j < 0$) and one with a positive value ($y_i > 0$). This is common when the measurand visualized by the bars does not match the measurand labeled on the bar graph's y-axis (usually differs by a log transformation). Similarly, consider the two lie factors normalized to different bars, and note the positive and negative quantities in the equation:

$$L_{i/j} = \frac{y_i - y_j}{y_j} \frac{x_j}{x_i - x_j} < 0, \tag{2.31}$$

$$L_{j/i} = \frac{y_i - y_j}{y_i} \frac{x_i}{x_i - x_j} > 0, \tag{2.32}$$

we get a positive and a negative lie factor normalized on different bars.

Note that for a well-represented graph, negative true values do not give a negative lie factor because the signs of the size of effects cancel.

How do we interpret negative lie factors? Once we found negative lie factors, the magnitude of the lie factors has less significance than the fact that they are negative. Having negative lie factors signifies that the figure is grossly misrepresented against basic visualization principles—the bars are either improperly baselined or do not represent the measurand as labeled on the axes. For quantification, figures containing negative lie factors should be removed from statistical calculations. However, they should be reported to quantify the extent of misrepresentation that falls into the above categories.

**2.2.1.5 Zero lie factors** Consider a misrepresented bar graph that has changes in true values ($x_i \neq x_j$) but shows no change in the visualized values ($y_i = y_j$). This may seem bizarre, but it is common for circular bar plots if the radius of each bar is carefully chosen. Extremely exaggerated bar plots that flood the visualized values to obscure the changes such that $y_i \approx y_j$ also satisfy such conditions. By definition, the lie factors are zero: $L_{i/j} = L_{j/i} = 0$.

**2.2.1.6 Undefined lie factors** Consider a misrepresented bar graph that has no change in true values ($x_i = x_j$) but shows changes in the visualized values ($y_i \neq y_j$). This may again seem bizarre, but it is common for circular bar plots, where the same true value has different visualized values depending on the radius of the bar. By definition, the lie factors are not defined (or infinity): $L_{i/j} = L_{j/i} = \infty$.

**2.2.1.7 Lie factors for multiple marks** Computing lie factors for graphs with two marks is straightforward. However, for graphs with multiple marks, one needs to consider all permutations of the pairwise comparisons between the marks, because the lie factors are defined such that they are normalized to one of the marks. This is justified because the audience has every right to select any pair of two marks and compare their differences based on one of the selected marks. For a graph with $n$ marks, we can define $m$ lie factors, where

$$m = P(n, 2) = \frac{n!}{(n-2)!} = n(n-1), \tag{2.33}$$

and $P$ denotes permutation. Consequently, the lie factor is not a graph-level property, but rather a property defined for permutations of two paired marks.

Interestingly, for graphs $n$ marks whose visualized value is linearly proportional to the true value, the number of effective lie factors is reduced to $n$ (instead of $m$) because the difference proportionality constant is a constant. For example, the lie factor normalized to mark $j$ comparing marks $i$ and $j$ becomes

$$L_{i/j} = \frac{\alpha_{i/j}}{\alpha_j} = \frac{k}{\alpha_j} \equiv L_j, \tag{2.34}$$

which is only dependent on mark $j$ used for normalization. Therefore, for linearly proportional relations between visualized and true values (not necessarily well-represented), the lie factor becomes a mark-level property, instead of a property defined for permutations of two paired marks.

##### 2.2.2 Steinbart's Graph Discrepancy Index (GDI)

In 1989, Paul Steinbart defined the graph discrepancy index (GDI) based on Tufte's lie factor to quantify data distortion in business annual reports [5]:

$$\mathrm{GDI}_{i/j} = 100(L_{i/j} - 1). \tag{2.35}$$

GDI has no practical difference compared to the lie factor. GDI is a mere change of range of the lie factor:

$$\begin{cases} \mathrm{GDI} \in (0, \infty) & \text{over-represented} \\ \mathrm{GDI} = 0 & \text{well-represented} \\ \mathrm{GDI} \in (-100, 0) & \text{under-represented} \end{cases} \tag{2.36}$$

for "well-behaved" graphs. The range of GDI for over- and under-represented graphs is similarly unbalanced, but it cannot be fixed by log transformation. GDI inherits the anomalous behavior of the lie factor when the graphs are not "well-behaved," [4] and the anomalous behavior is harder to detect because of the shift from the lie factor from GDI's definition. GDI gained wide usage in the business literature, but it should be refrained from further usage due to such disadvantages.

##### 2.2.3 Lie Factor of Fold Change

**2.2.3.1 Mathematical definition of the lie factors of fold change** We can similarly define a **lie factor of fold change** as

$$\text{Lie factor of fold change} = \frac{\text{Fold change of visualized value}}{\text{Fold change of true value}}, \tag{2.37}$$

where the fold change is defined by

$$\text{Fold change} = \frac{\text{Value 2}}{\text{Value 1}}. \tag{2.38}$$

Mathematically, we can define the lie factor of the fold change between mark $i$ and mark $j$'s magnitude channel as

$$F_{i/j} \equiv \frac{y_i/y_j}{x_i/x_j} = \frac{y_i x_j}{y_j x_i}. \tag{2.39}$$

Simplify by definition of mark proportionality constant and Equation 2.23, we have

$$F_{i/j} = \frac{\alpha_i}{\alpha_j} = \frac{L_{i/j}}{L_{j/i}}, \tag{2.40}$$

suggesting that the lie factor of fold change is the ratio between mark proportionality constants of marks $i$ and $j$, which is equivalent to the ratio between lie factors normalized to marks $j$ and $i$.

**2.2.3.2 Lie factors of fold change for multiple marks** By Equation 2.40, the lie factors of fold change for marks $i$ and $j$ follow an inverse relation:

$$F_{i/j} = F_{j/i}^{-1}, \tag{2.41}$$

suggesting that they are not independent of each other. Lie factors of fold change is therefore a useful metric on a pairwise-combination level. When quantifying $F_{i/j}$, one can arbitrarily choose the order of the $i$-$j$ combination such that $|x_i| > |x_j|$ or $|x_i| < |x_j|$, but such choice must be reported. For a graph with $n$ marks, we can define $m$ lie factors of fold change, where

$$m = C(n, 2) = \frac{n!}{2!(n-2)!} = \frac{n}{2}(n-1), \tag{2.42}$$

and $C$ denotes combination.

**2.2.3.3 Behavior of the lie factors of fold change** If the graph is well-represented, the mark proportionality constants should be the same, so the lie factors of fold change simplify to unity:

$$F_{i/j} = \frac{\alpha_i}{\alpha_j} = 1 \tag{2.43}$$

$$F_{j/i} = \frac{\alpha_j}{\alpha_i} = 1 \tag{2.44}$$

What happens when the graph is misrepresented? Consider $F_{i/j}$ and $i$-$j$ combinations that satisfy $|x_i| > |x_j|$. If the visualized values over-represent the true value, the lie factors are greater than 1; if the visualized values under-represent the true value, the lie factors are positive and less than 1. Let's consider a representative example for analysis.

For the following analysis, consider misrepresented but "well-behaved" graphs that (1) have positive values ($x_i > 0, x_j > 0$) and (2) do not visualize positive values with negative magnitude channels and vice versa ($\text{sgn}(x_i) = \text{sgn}(y_i)$, $\text{sgn}(x_j) = \text{sgn}(y_j)$). Consider the case of $x_i > x_j > 0$, $y_i > y_j > 0$. To over-represent the true value, either $y_j$ decreases by $k_j > 0$ or $y_i$ increases by $k_i > 0$ in the over-represented lie factor of fold change $F'_{i/j}$:

$$\begin{aligned} F'_{i/j} &= \frac{y'_i \uparrow x_j}{y'_j \downarrow x_i} \\ &= \frac{y_i + k_i}{y_j - k_j} \frac{x_j}{x_i} \end{aligned}$$

We ask if the over-represented lie factor of fold change is greater than the well-represented one $F_{i/j} = 1$:

$$\begin{aligned} F'_{i/j} &\overset{?}{>} F_{i/j} \\ \frac{y_i + k_i}{y_j - k_j} \frac{x_j}{x_i} &\overset{?}{>} \frac{y_i}{y_j} \frac{x_j}{x_i} \\ \frac{y_i + k_i}{y_j - k_j} &\overset{?}{>} \frac{y_i}{y_j} \\ y_i y_j + k_i y_j &\overset{?}{>} y_i y_j - k_j y_i \\ k_i y_j &> -k_j y_i \end{aligned} \qquad \blacksquare$$

Because all the quantities are positive, we have proved that $F'_{i/j} > 1$ subjected to the above assumptions. The result is also intuitive to understand from the perspective of mark proportionality constants. To over-represent the true value, either $\alpha_i$ increases and/or $\alpha_j$ decreases, which all results in $F'_{i/j} > F_{i/j} = 1$. Similar logic applies to "well-behaved" graphs with negative values ($x_i < 0, x_j < 0$). Full proof for all cases is provided in Supplementary Proofs.

**2.2.3.4 Interpretation of the lie factors of fold change** In summary, for "well-behaved" graphs, $F_{i/j}$ subjected to $|x_i| > |x_j|$ satisfies

$$\begin{cases} F_{i/j} \in (1, \infty) & \text{over-represented} \\ F_{i/j} = 1 & \text{well-represented} \\ F_{i/j} \in (0, 1) & \text{under-represented} \end{cases} \tag{2.45}$$

Like the lie factor of relative change, the lie factor of fold change's over-representation and under-representation of graphs have different range scales. This can be fixed by using the log-transformed lie factor of fold change:

$$\begin{cases} \log(F_{i/j}) \in (0, \infty) & \text{over-represented} \\ \log(F_{i/j}) = 0 & \text{well-represented} \\ \log(F_{i/j}) \in (-\infty, 0) & \text{under-represented} \end{cases} \tag{2.46}$$

A form of $\log(F_{i/j})$ is mentioned in the footnote of prior research [4] as the log discrepancy index, but it is never explored or characterized.

By rearranging the lie factor of fold change equation, we have five interpretations for the lie factor of fold change. The lie factor of fold change quantifies the ratio between the fold change in visual value and fold change in true value:

$$F_{i/j} \equiv \frac{y_i/y_j}{x_i/x_j}. \tag{2.39}$$

The lie factor of fold change between marks $i$ and $j$ is the ratio between the mark proportionality constants of marks $i$ and $j$:

$$F_{i/j} = \frac{\alpha_i}{\alpha_j}. \tag{2.21}$$

The lie factor of fold change is also the ratio between the lie factor of relative change of mark $i$ normalized to mark $j$ and lie factor of relative change of mark $j$ normalized to mark $i$:

$$F_{i/j} = \frac{L_{i/j}}{L_{j/i}}. \tag{2.21}$$

Rearranging Equation 2.39, the lie factor of fold change can be expressed as the fold change of visualized value of mark $i$ compared to the theoretical visualized value of mark $i$ based on the mark proportionality constant of mark $j$:

$$F_{i/j} = \frac{y_i}{y'_{i/j}}. \tag{2.47}$$

The lie factor of fold change can also be rearranged to the fold change of the theoretical visualized value of mark $j$ based on the mark proportionality constant of mark $i$ compared to the visualized value of mark $j$:

$$F_{i/j} = \frac{y'_{j/i}}{y_j}. \tag{2.48}$$

Note that because the lie factor of fold change is based on fold change of absolute magnitude, it does not reflect the relative magnitude of the change.

**2.2.3.5 Negative lie factors of fold change** The preceding analysis covers "well-behaved" graphs that we encounter most of the time. However, it breaks down when graphs are misrepresented and not "well-behaved." In this section, consider misrepresented and not "well-behaved" graphs that may give rise to negative lie factors of fold change.

Similar to lie factors of relative change, negative lie factor of fold change can are common for bar graphs whose direction carries information about the value's sign. If one of the mark proportionality constants is negative (i.e. a visualized value has a different sign than its true value), the lie factor of fold change is negative. Two examples illustrated in Section 2.2.1.4 apply similarly to lie factors of fold change.

Note that if both marks used in a lie factor of fold change have such sign mismatch, the lie factor of fold change is positive, causing such anomaly not easily detected. Therefore, if a graph has a negative lie factor of fold change, the graph's design must be inspected with caution to interpret other lie factors of fold change.

Like lie factors of relative change, having negative lie factors of fold change signifies that the figure is grossly misrepresented against basic visualization principles–the bars are either improperly baselined or do not represent the measurand as labeled on the axes. For quantification, figures containing negative lie factors of fold change should be removed from statistical calculations. However, they should be reported to quantify the extent of misrepresentation that falls into the above categories.

**2.2.3.6 Zero or undefined lie factors of fold change** The lie factor of fold change is well-defined for graphs that satisfy $x_i \neq x_j$ or $y_i \neq y_j$, therefore those conditions do not cause anomalous lie factor of fold change. To have zero or undefined lie factor of fold change, mark proportionality constants in the numerator or denominator of Equation 2.39 must be zero, which applies to almost no practical cases.

##### 2.2.4 Mather's Relative Graph Discrepancy (RGD)

In search for an alternative metric to quantify data distortion other than the lie factor or GDI, Manther in 2005 defined the relative graph discrepancy (RGD) [4] of mark $i$ given the mark proportionality constant of mark $j$ as

$$\text{RGD}_{i/j} = \frac{y_i - y'_{i/j}}{y'_{i/j}}, \tag{2.49}$$

which quantifies the relative deviation of the visualized value of mark $i$ compared to its theoretical visualized value given $\alpha_j$. Rearrange, we also obtain

$$\text{RGD}_{i/j} = \frac{y'_{j/i} - y_j}{y_j}, \tag{2.50}$$

which gives $\text{RGD}_{i/j}$ the interpretation of the relative deviation of the theoretical visualized value of mark $j$ given $\alpha_i$ compared to the visualized value of mark $i$. We can also simplify Equation 2.49 to get

$$\text{RGD}_{i/j} = F_{i/j} - 1, \tag{2.51}$$

suggesting $\text{RGD}_{i/j}$ is a simple range transformation of $F_{i/j}$, which allows $\text{RGD}_{i/j}$ to inherit most of the properties of $F_{i/j}$.

For "well-behaved" graphs, $\text{RGD}_{i/j}$ subjected to $|y_i| > |y_j|$ satisfies

$$\begin{cases} \text{RGD}_{i/j} \in (0, \infty) & \text{under-represented} \\ \text{RGD}_{i/j} = 0 & \text{well-represented} \\ \text{RGD}_{i/j} \in (-1, 0) & \text{over-represented} \end{cases} \tag{2.52}$$

The range of $\text{RGD}_{i/j}$ is similarly unbalanced, and it cannot be fixed by log transformation. $\text{RGD}_{i/j}$ also inherits the anomalous behavior of $F_{i/j}$ when the graphs are not "well-behaved." Because of the additional disadvantages of $\text{RGD}_{i/j}$ compared to $F_{i/j}$, $\text{RGD}_{i/j}$ should be refrained from further usage.

##### 2.2.5 Normalized Variability Metrics

By the principle of proportional ink, the mark proportionality constants of all marks should be the same for a well-represented graph but not necessarily for a misrepresented graph. We can therefore quantify the extent of data distortion using the extent of variability of mark proportionality constants. Similar concept applies to difference proportionality constants but with some caveats. We explore normalized variability metrics of coefficient of variation and its robust version based on interquartile range and median absolute deviation.

**2.2.5.1 Coefficient of variation (CV)** The non-dimensional **coefficient of variation (CV) of mark proportionality constants** is defined as

$$\mathrm{CV}_{\alpha_i} = \frac{\sigma_{\alpha_i}}{\mu_{\alpha_i}}, \tag{2.53}$$

where $\sigma_{\alpha_i}$ is the standard deviation of mark proportional constants, and $\mu_{\alpha_i}$ is the mean of mark proportional constants. Note that $\mathrm{CV}_{\alpha_i}$ is a summary statistic of the population of mark proportionality constants of all marks of a graph, therefore being a graph-level statistic.

$\mathrm{CV}_{\alpha_i}$ inherits all properties of CV as defined in statistics. When there is no variation in mark proportionality constant (i.e. all the same in a well-represented graph), $\mathrm{CV}_{\alpha_i} = 0$. For a misrepresented graph, the mark proportionality constants are not the same, having $\mathrm{CV}_{\alpha_i} > 0$. In summary,

$$\mathrm{CV}_{\alpha_i} \begin{cases} = 0 & \text{well-represented} \\ > 0 & \text{misrepresented} \end{cases}. \tag{2.54}$$

Note that $\mathrm{CV}_{\alpha_i}$ does not distinguish over- and under-representations. $\mathrm{CV}_{\alpha_i}$ quantifies the extent of variation in the mark proportionality constant: the standard deviation of mark proportionality constants normalized to their mean.

We can similarly define the **coefficient of variation of difference proportionality constant** as

$$\mathrm{CV}_{\alpha_{i/j}} = \frac{\sigma_{\alpha_{i/j}}}{\mu_{\alpha_{i/j}}}, \tag{2.55}$$

where $\sigma_{\alpha_{i/j}}$ is the standard deviation of difference proportional constants, and $\mu_{\alpha_{i/j}}$ is the mean of difference proportional constants. Note that $\mathrm{CV}_{\alpha_{i/j}}$ is best used for misrepresented graphs that have a nonlinear relationship between visualized value and true value because $\alpha_{i/j}$ is a constant for linearly proportional relationships ($\mathrm{CV}_{\alpha_{i/j}} = 0$, Section 2.1.2.3). Therefore,

$$\mathrm{CV}_{\alpha_{i/j}} \begin{cases} = 0 & \text{linearly proportional representation} \\ > 0 & \text{misrepresented} \end{cases}. \tag{2.56}$$

Note that linearly proportional representation could be well-represented (directly proportional, $b = 0$) or misrepresented ($b \neq 0$).

Because CV is based on standard deviation and mean, it is prone to influence by outliers in the data. To improve the robustness of normalized variability metrics, we can define more robust versions of CV based on interquartile range and median absolute deviations [6], explored below.

**2.2.5.2 Robust CV based on interquartile range (RCVQ)** Interquartile range (IQR) is a robust measure of variability, defined as the difference between the sample's 75 percentile and 25 percentile. We can define the **robust coefficient of variation based on IQR** (RCVQ) as

$$\mathrm{RCVQ} = 0.75\frac{\mathrm{IQR}}{\mathrm{median}}, \tag{2.57}$$

where 0.75 is the consistency factor for equivalent comparisons with CV for a normal distribution [6]. We can use RCVQ for both mark proportionality constants ($\mathrm{RCVQ}_{\alpha_i}$) and difference proportionality constants ($\mathrm{RCVQ}_{\alpha_i/j}$). Similar with CV, we can interpret RCVQs with:

$$\mathrm{RCVQ}_{\alpha_i} \begin{cases} = 0 & \text{well-represented} \\ > 0 & \text{misrepresented} \end{cases} \tag{2.58}$$

and

$$\mathrm{RCVQ}_{\alpha_{i/j}} \begin{cases} = 0 & \text{linearly proportional representation} \\ > 0 & \text{misrepresented} \end{cases}. \tag{2.59}$$

**2.2.5.3 Robust CV based on median absolute deviation (RCVM)** Another robust measure of variability is median absolute deviation (MAD), defined as the median of the set of absolute deviations of each element in the population with the median of the population:

$$\text{MAD} = \text{median}(|x_i - \text{median}(\mathbf{x})|), \tag{2.60}$$

where $x_i$ is an element in the population, and $\mathbf{x}$ is the set of all elements in the population. We can define the **robust coefficient of variation based on MAD** (RCVM) as

$$\text{RCVM} = 1.4826 \frac{\text{MAD}}{\text{median}}, \tag{2.61}$$

where 1.4826 is the consistency factor for equivalent comparisons with CV for a normal distribution [6]. We can use RCVM for both mark proportionality constants ($\text{RCVM}_{\alpha_i}$) and difference proportionality constants ($\text{RCVM}_{\alpha_{i/j}}$). Similar with CV, we can interpret RCVMs with:

$$\text{RCVM}_{\alpha_i} \begin{cases} = 0 & \text{well-represented} \\ > 0 & \text{misrepresented} \end{cases} \tag{2.62}$$

and

$$\text{RCVM}_{\alpha_{i/j}} \begin{cases} = 0 & \text{linearly proportional representation} \\ > 0 & \text{misrepresented} \end{cases}. \tag{2.63}$$

**2.2.5.4 Normalized variability metrics for zeroing mistakes** Because the different proportionality constant is the same for linearly proportional and directly proportional relationships between visualized and true values (Eq 2.15)

$$\alpha_{i/j} = k \equiv \alpha, \tag{2.15}$$

we have a ground truth for the graph proportionality constant for a bar graph with zeroing mistakes. Therefore, instead of normalizing to the mean or median of the mark proportionality constants (a guess of the constant's groundtruth), we can normalize directly to the groundtruth of graph proportionality constant, which is equivalent to the difference proportionality constant for graphs with zeroing mistakes.

All of the above metrics can be rewritten for zeroing mistakes with the following definitions, where the variables are similar defined as above. The CV for zeroing mistakes (CVZ) of mark proportionality constant can be defined as

$$\text{CVZ}_{\alpha_i} = \frac{\sigma_{\alpha_i}}{\alpha}. \tag{2.64}$$

The RCVQ for zeroing mistakes (RCVQZ) of mark proportionality constant can be defined as

$$\text{RCVQZ} = 0.75 \frac{\text{IQR}}{\alpha}. \tag{2.65}$$

The RCVM for zeroing mistakes (RCVMZ) of mark proportionality constant can be defined as

$$\text{RCVMZ} = 1.4826 \frac{\text{MAD}}{\alpha}. \tag{2.66}$$

Note that these definitions only apply to bar graphs with zeroing mistakes but not log mistakes. Graphs with log mistakes does not have similar properties because there does not exist a groundtruth proportionality constant (i.e. bar graphs are not meant for y-axis in log scale).

##### 2.2.6 Mitigating Data Biases

When using the data distortion metrics, we need to be cautious of (1) **bar-level biases** caused by the uneven distribution of the number of bars in each graph and (2) **graph-level biases** caused by the uneven distribution of the number of graphs in each journal article. Bar-level bias needs to be mitigated for bar-level metrics that are based on pairwise-permutation or pairwise-combination of bars. Graph-level bias needs to be mitigated for bar-level metrics after bar-level bias is mitigated and graph-level metrics.

As an example for bar-level bias, suppose an article has 7 misused bar graphs, and they each have {14, 14, 8, 2, 2, 2, 2} quantifiable bars. Because lie factor of relative change is a pairwise-permutation level metric, each graph can generate {91, 91, 28, 2, 2, 2, 2} lie factors. Therefore, the graph with more bars will dominate the dataset, and the effect of graphs with less bars will be negligible. To mitigate such bar-level data bias, we can take an ensemble approach by taking the median lie factor of each graph. After the correction, we have one lie factor for each graph.

Graph-level bias are common because articles nowadays are able to perform similar experiments on different samples, and similar graphs can be shown in supplementary information. As an example for graph-level bias, suppose we 3 articles with different numbers and measurand annotations of misused graphs: (A) 7 graphs representing "percentage," (B) 1 graph representing "ratio" and 1 graph representing "concentration," and (C) 2 graphs representing "count," 1 graph representing "time," and 5 graphs representing "percentage." Each graph has a median lie factor of relative change, but the lie factor dataset is dominated by articles A and C because they have a lot more graphs than article B. To mitigate such graph-level data bias, we can again take an ensemble approach by grouping graphs within the sample article according to their measurand annotation and take the median of their lie factors. After correction, we have 1 median lie factor representing each measurand annotation within each graph: (A) 1 median lie factor representing "percentage" graphs, (B) 1 median lie factor representing "ratio" graphs and 1 median lie factor representing "concentration" graphs, (C) 1 median lie factor representing "count" graphs, 1 median lie factor representing "time" graphs, and 1 median lie factor representing "percentage" graphs. This approach avoids the graphs with the same measurand spamming the dataset.

#### 2.3 Bar Graphs in Polar Coordinates

Bar graphs in polar coordinates are usually used to display a large number of data and are not as commonly used as those in Cartesian coordinates. However, we show that the bar graphs in polar coordinates inherently distorts the underlying data and should not be used for displaying scientific data.

##### 2.3.1 Radial Bar Graphs

**2.3.1.1 Violation of Principle of Proportional Ink** Radial bar graphs are bar graphs in the polar coordinates, where the bars are arranged categorically along the radial coordinate, and the true value is encoded along the angular coordinate. Radial bar graphs intend to make the angle of each bar the magnitude channel, but the area of each bar is not directly proportional to the angle. In the following section, we prove the failure of direct proportionality between the area of each bar and the true value of radial bar graphs.

Consider a radial bar graph with $n$ bars, each with a constant width $W$ arranged radially. The inner edge of the $i$th bar has a radius $r_i$, and the outer edge of the $i$ th bar has a radius $R_i \equiv r_i + W$. The "height" of each bar is directly proportional to the angular coordinate $\theta_i$. We ask if the area of each bar $A_i$ is a proportional function of $\theta_i$ with proportionality constant $\beta_i$:

$$A_i = \beta_i \theta_i. \tag{2.67}$$

The area of each bar can be calculated using the area formula of sectors:

$$\begin{aligned}A_i &= \frac{\theta_i}{2\pi}\pi(R_i^2 - r_i^2) \\ &= \frac{\theta_i}{2}((r_i + W)^2 - r_i^2) \\ &= \frac{\theta_i}{2}(r_i^2 + 2Wr_i + W^2 - r_i^2) \\ &= \frac{W\theta_i}{2}(2r_i + W),\end{aligned}$$

which shows that $A_i = f(r_i, \theta_i)$ and $\beta_i(r_i) = Wr_i + \frac{1}{2}W^2$. The derivation shows that the area of each bar is arbitrarily dependent on the radial coordinate, which is related to the categorical variable, not the true value, therefore violating the principle of proportional ink.

**2.3.1.2 Quantifying Data Distortion** The lie factor of bars $i$ and $j$ in a radial bar plot is

$$\begin{aligned}L_{i/j} &= \frac{A_i - A_j}{A_j}\frac{\theta_j}{\theta_i - \theta_j} \\ &= \frac{\theta_i(2r_i + W) - \theta_j(2r_j + W)}{\theta_j(2r_j + W)}\frac{\theta_j}{\theta_i - \theta_j} \\ &= \frac{\theta_i(2r_i + W) - \theta_j(2r_j + W)}{\theta_i(2r_j + W) - \theta_j(2r_j + W)}\end{aligned}$$

The lie factor of fold change of bars $i$ and $j$ in a radial bar plot is

$$F_{i/j} = \frac{A_i}{A_j}\frac{\theta_j}{\theta_i} \tag{2.68}$$

$$= \frac{\theta_i(2r_i + W)}{\theta_j(2r_j + W)}\frac{\theta_j}{\theta_i} \tag{2.69}$$

$$= \frac{2r_i + W}{2r_j + W} \tag{2.70}$$

The coefficient of variation of mark proportionality constant in a radial bar plot is

$$\mathrm{CV}_{\beta_i} = \mathrm{CV}\left(Wr_i + \frac{1}{2}W^2\right)$$

##### 2.3.2 Circular Bar Graphs

**2.3.2.1 Violation of Principle of Proportional Ink** Circular bar graphs are bar graphs in the polar coordinates, where the bars are arranged categorically along the angular coordinate, and the true value is encoded along the radial coordinate. Angular bar graphs intend to make the height of each bar the magnitude channel, but the area of each bar is not directly proportional to the height. In the following section, we prove the failure of direct proportionality between the area of each bar and the true value of circular bar graphs.

Consider a circular bar graph with $n$ bars, each occupying a constant angle $\theta$ arranged angularly. The inner edge of each bar has the same radius $r$. The outer edge of the $i$th bar has a radius $R_i$, which is related to the height of each bar $H_i$ by $R_i \equiv r + H_i$. We ask if the area of each bar $A_i$ is directly proportional to $H_i$ with proportionality constant $\beta_i$:

$$A_i = \beta_i H_i \tag{2.71}$$

The area of each bar can be calculated using the area formula of sectors:

$$\begin{aligned} A_i &= \frac{\theta}{2\pi}\pi(R_i^2 - r^2) \\ &= \frac{\theta}{2}((r + H_i)^2 - r^2) \\ &= \frac{\theta}{2}(r^2 + 2rH_i + H_i^2 - r^2) \\ &= \frac{\theta}{2}(H_i^2 + 2rH_i), \end{aligned}$$

which shows that $A_i$ depends quadratically on $H_i$, and $\beta_i(H_i) = \frac{1}{2}\theta H_i + \theta r$. Therefore, the area of each bar is not directly proportional to the true values, violating the principle of proportional ink.

A polar area chart is a simplified circular bar graph where the inner radius $r = 0$, which also violates the principle of proportional ink.

**2.3.2.2 Quantifying Data Distortion** The lie factor of bars $i$ and $j$ in a circular bar plot is

$$\begin{aligned} L_{i/j} &= \frac{A_i - A_j}{A_j}\frac{H_j}{H_i - H_j} \\ &= \frac{H_i^2 + 2rH_i - H_j^2 - 2rH_j}{H_j^2 + 2rH_j}\frac{H_j}{H_i - H_j} \\ &= \frac{(H_i^2 - H_j^2) + 2r(H_i - H_j)}{H_j + 2r}\frac{1}{H_i - H_j} \\ &= \frac{(H_i + H_j + 2r)(H_i - H_j)}{H_j + 2r}\frac{1}{H_i - H_j} \\ &= \frac{H_i + H_j + 2r}{H_j + 2r} \end{aligned}$$

The lie factor of fold change of bars $i$ and $j$ in a circular bar plot is

$$F_{i/j} = \frac{A_i}{A_j}\frac{H_j}{H_i} \tag{2.72}$$

$$= \frac{H_i^2 + 2rH_i}{H_j^2 + 2rH_j}\frac{H_j}{H_i} \tag{2.73}$$

$$= \frac{H_i + 2r}{H_j + 2r} \tag{2.74}$$

The coefficient of variation of mark proportionality constant in a circular bar plot is

$$\mathrm{CV}_{\beta_i} = \mathrm{CV}\left(\frac{1}{2}\theta H_i + \theta r\right)$$

#### 2.4 Preventing Visualization Mistakes

To prevent visualization mistakes, hierarchy of controls need to be implemented. Here, we discuss details about our proposed engineering controls on software and administrative controls of having journal visualization checklists for authors, reviewers, and editors.

##### 2.4.1 Software engineering controls

Plotting packages in programming languages allows maximal freedom in graph creation, but it comes with the risks of creating graphs that do not follow the principle of proportional ink. For example, the Matplotlib package [7] in Python not only enables the creation of bar plots with log mistakes without warning, but also has tutorials actively teaching people how to create them [8]. Matplotlib also has commands to change the bars' baseline [9] and does not have software warnings to prevent people from modifying the baseline of the bars. Similar issues occur in seaborn in Python [10], ggplot in R [11], and MATLAB®[12]. Although the documentation of ggplot warns programmers of the risk of changing the axis scaling of bar graphs [13], the warning is buried in technical details and not accessible to programmers when mistakes are being made.

We propose that plotting packages in programming languages should have built-in software-level warnings that notify the programmers of visualization mistakes made by axis distortion. For example, when attempting to change the axis scaling to logarithmic or change the baseline to start at 0, a warning message should be printed to the software log or graphic user interface.

Specialized statistics and graphing software are tailored to audiences with less programming experience and require more guidance in creating statistical analyses and data visualizations. However, software like GraphPad Prism [14] provides the users the option to change the axis scaling to logarithmic and to change the baseline of the bars without issuing warnings. Although GraphPad Prism provides warnings against log mistakes in bar graphs in its documentation [15] and FAQ sections [16, 17], those warnings are hard to locate unless directly searched on the web. Additionally, no warnings are issued when those mistakes are made in the software.

We propose that statistics and graphing software should not only have more detailed warnings, guidelines, and visualization suggestions for accurate data visualization in the documentation, but also have software-level warnings when users are selecting options that may distort the underlying data. For example, when the user clicks on the option to set the bar's baseline to a nonzero value, warnings need to be displayed in the interface to notify the consequences of making such adjustments.

##### 2.4.2 Journal visualization checklists

In addition to the current journal visualization checklists that focus on data transparency (i.e. individual data points shown whenever possible, data distribution shown whenever possible), we propose an additional checklist for the correct use of bar graphs, especially for biological research articles. For each bar graph, do they

- have Cartesian coordinates, but not polar coordinates,
- have a linear y-axis (axis for the numerical variable) that starts at 0,
- have $\geq 2$ bars,
- have the same bar width for all bars,
- have y-axis tick label for 0,
- have bars in two dimension (2D), but not three dimension (3D),
- have a measurand that has defined relative difference, and
- have raw data shown.

For each bar graph with discontinuous (broken) axis, do they satisfy the additional guidelines of

- explicitly break both the axis and the bars,
- have $\geq 2$ bars below the break,
- have y-axis tick labels for numerical values at the break, and

- have no stacks.

For each stacked bar graph, do they satisfy the additional guidelines of

- have no breaks, and
- have $\leq 2$ stacks for general stacked bar graphs or $\leq 3$ stacks for percent stacked bar graphs.

#### 2.5 Supplementary Proofs

##### 2.5.1 Derivation of lie factor for positive values

Consider misrepresented but "well-behaved" graphs that (1) have positive values ($x_i > 0, x_j > 0$) and (2) do not visualize positive values with negative magnitude channels and vice versa ($\text{sgn}(x_i) = \text{sgn}(y_i)$, $\text{sgn}(x_j) = \text{sgn}(y_j)$).

**Case 1: Over-representation of $L'_{i/j}$, $x_i > x_j > 0$, $y_i > y_j > 0$.** To over-represent the true value, either $y_j$ decreases by $k_j > 0$ or $y_i$ increases by $k_i > 0$ in the over-represented lie factor $L'_{i/j}$:

$$\begin{aligned} L'_{i/j} &= \frac{y'_i \uparrow - y'_j \downarrow}{y'_j \downarrow} \frac{x_j}{x_i - x_j} \\ &= \frac{(y_i + k_i) - (y_j - k_j)}{y_j - k_j} \frac{x_j}{x_i - x_j} \end{aligned}$$

We ask if the over-represented lie factor is greater than the well-represented lie factor $L_{i/j} = 1$:

$$\begin{aligned} L'_{i/j} &\stackrel{?}{>} L_{i/j} \\ \frac{(y_i + k_i) - (y_j - k_j)}{y_j - k_j} \frac{x_j}{x_i - x_j} &\stackrel{?}{>} \frac{y_i - y_j}{y_j} \frac{x_j}{x_i - x_j} \\ \frac{y_i - y_j + k_i + k_j}{y_j - k_j} &\stackrel{?}{>} \frac{y_i - y_j}{y_j} \\ y_i y_j - y_j^2 + k_i y_j + k_j y_j &\stackrel{?}{>} y_i y_j - y_j^2 - y_i k_j + y_j k_j \\ k_i y_j &> -y_i k_j \qquad \blacksquare \end{aligned}$$

Therefore, $L'_{i/j} > L_{i/j} = 1$.

**Case 2: Over-representation of $L'_{j/i}$, $x_i > x_j > 0$, $y_i > y_j > 0$.** To over-represent the true value, either $y_j$ decreases by $k_j > 0$ or $y_i$ increases by $k_i > 0$ in the over-represented lie factor $L'_{j/i}$:

$$\begin{aligned} L'_{j/i} &= \frac{y'_i \uparrow - y'_j \downarrow}{y'_i \uparrow} \frac{x_j}{x_i - x_j} \\ &= \frac{(y_i + k_i) - (y_j - k_j)}{y_i + k_i} \frac{x_j}{x_i - x_j} \end{aligned}$$

We ask if the over-represented lie factor is greater than the well-represented lie factor $L_{j/i} = 1$:

$$\begin{aligned} L'_{j/i} &\stackrel{?}{>} L_{j/i} \\ \frac{(y_i + k_i) - (y_j - k_j)}{y_i + k_i} \frac{x_j}{x_i - x_j} &\stackrel{?}{>} \frac{y_i - y_j}{y_i} \frac{x_j}{x_i - x_j} \\ \frac{y_i - y_j + k_i + k_j}{y_i + k_i} &\stackrel{?}{>} \frac{y_i - y_j}{y_i} \\ y_i^2 - y_i y_j + k_i y_i + k_j y_i &\stackrel{?}{>} y_i^2 + k_i y_i - y_i y_j - y_j k_i \\ k_j y_i &> -y_j k_i \qquad \blacksquare \end{aligned}$$

Therefore, $L'_{j/i} > L_{j/i} = 1$.

**Case 3: Over-representation of $L'_{i/j}$, $x_j > x_i > 0$, $y_j > y_i > 0$.** Swap index such that $i \leftarrow j$ and $j \leftarrow i$. The proof is identical to case 2. Therefore, $L'_{i/j} > L_{i/j} = 1$.

**Case 4: Over-representation of $L'_{j/i}$, $x_j > x_i > 0$, $y_j > y_i > 0$.** Swap index such that $i \leftarrow j$ and $j \leftarrow i$. The proof is identical to case 1. Therefore, $L'_{j/i} > L_{j/i} = 1$.

**Case 5: Under-representation of $L'_{i/j}$, $x_i > x_j > 0$, $y_i > y_j > 0$.** To under-represent the true value, either $y_j$ increases by $k_j > 0$ or $y_i$ decreases by $k_i > 0$ in the under-represented lie factor $L'_{i/j}$:

$$\begin{aligned} L'_{i/j} &= \frac{y'_i \downarrow - y'_j \uparrow}{y'_j \uparrow} \frac{x_j}{x_i - x_j} \\ &= \frac{(y_i - k_i) - (y_j + k_j)}{y_j + k_j} \frac{x_j}{x_i - x_j} \end{aligned}$$

We ask if the under-represented lie factor is less than the well-represented lie factor $L_{i/j} = 1$:

$$\begin{aligned} L'_{i/j} &\overset{?}{<} L_{i/j} \\ \frac{(y_i - k_i) - (y_j + k_j)}{y_j + k_j} \frac{x_j}{x_i - x_j} &\overset{?}{<} \frac{y_i - y_j}{y_j} \frac{x_j}{x_i - x_j} \\ \frac{y_i - y_j - k_i - k_j}{y_j + k_j} &\overset{?}{<} \frac{y_i - y_j}{y_j} \\ y_i y_j - y_j^2 - k_i y_j - k_j y_j &\overset{?}{<} y_i y_j - y_j^2 + y_i k_j - y_j k_j \\ -k_i y_j &< y_i k_j \end{aligned} \quad \blacksquare$$

Therefore, $L'_{i/j} < L_{i/j} = 1$.

**Case 6: Under-representation of $L'_{j/i}$, $x_i > x_j > 0$, $y_i > y_j > 0$.** To under-represent the true value, either $y_j$ increases by $k_j > 0$ or $y_i$ decreases by $k_i > 0$ in the under-represented lie factor $L'_{j/i}$:

$$\begin{aligned} L'_{j/i} &= \frac{y'_i \downarrow - y'_j \uparrow}{y'_i \downarrow} \frac{x_j}{x_i - x_j} \\ &= \frac{(y_i - k_i) - (y_j + k_j)}{y_i - k_i} \frac{x_j}{x_i - x_j} \end{aligned}$$

We ask if the under-represented lie factor is less than the well-represented lie factor $L_{j/i} = 1$:

$$\begin{aligned} L'_{j/i} &\overset{?}{<} L_{j/i} \\ \frac{(y_i - k_i) - (y_j + k_j)}{y_i - k_i} \frac{x_j}{x_i - x_j} &\overset{?}{<} \frac{y_i - y_j}{y_i} \frac{x_j}{x_i - x_j} \\ \frac{y_i - y_j - k_i - k_j}{y_i - k_i} &\overset{?}{<} \frac{y_i - y_j}{y_i} \\ y_i^2 - y_i y_j - k_i y_i - k_j y_i &\overset{?}{<} y_i^2 - k_i y_i - y_i y_j + y_j k_i \\ -k_j y_i &< y_j k_i \end{aligned} \quad \blacksquare$$

Therefore, $L'_{j/i} < L_{j/i} = 1$.

**Case 7: Under-representation of $L'_{i/j}$, $x_j > x_i > 0$, $y_j > y_i > 0$.** Swap index such that $i \leftarrow j$ and $j \leftarrow i$. The proof is identical to case 6. Therefore, $L'_{i/j} < L_{i/j} = 1$.

**Case 8: Under-representation of $L'_{j/i}$, $x_j > x_i > 0$, $y_j > y_i > 0$.** Swap index such that $i \leftarrow j$ and $j \leftarrow i$. The proof is identical to case 5. Therefore, $L'_{j/i} < L_{j/i} = 1$.

##### 2.5.2 Derivation of lie factor for negative values

Consider misrepresented but "well-behaved" graphs that (1) have negative values ($x_i < 0, x_j < 0$) and (2) do not visualize positive values with negative magnitude channels and vice versa ($\text{sgn}(x_i) = \text{sgn}(y_i)$, $\text{sgn}(x_j) = \text{sgn}(y_j)$).

**Case 1: Over-representation of $L'_{i/j}$, $x_i < x_j < 0$, $y_i < y_j < 0$.** To over-represent the true value, either $y_j$ increase by $k_j > 0$ or $y_i$ decrease by $k_i > 0$ in the over-represented lie factor $L'_{i/j}$:

$$
\begin{aligned}
L'_{i/j} &= \frac{y'_i \downarrow - y'_j \uparrow}{y'_j \uparrow} \frac{x_j}{x_i - x_j} \\
&= \frac{(y_i - k_i) - (y_j + k_j)}{y_j + k_j} \frac{x_j}{x_i - x_j}
\end{aligned}
$$

We ask if the over-represented lie factor is greater than the well-represented lie factor $L_{i/j} = 1$:

$$
\begin{aligned}
L'_{i/j} &\overset{?}{>} L_{i/j} \\
\frac{(y_i - k_i) - (y_j + k_j)}{y_j + k_j} \frac{x_j}{x_i - x_j} &\overset{?}{>} \frac{y_i - y_j}{y_j} \frac{x_j}{x_i - x_j} \\
\frac{y_i - y_j - k_i - k_j}{y_j + k_j} &\overset{?}{>} \frac{y_i - y_j}{y_j} \\
y_i y_j - y_j^2 - k_i y_j - k_j y_j &\overset{?}{>} y_i y_j - y_j^2 + y_i k_j - y_j k_j \\
-k_i y_j &> y_i k_j \qquad \blacksquare
\end{aligned}
$$

Therefore, $L'_{i/j} > L_{i/j} = 1$.

**Case 2: Over-representation of $L'_{j/i}$, $x_i < x_j < 0$, $y_i < y_j < 0$.** To over-represent the true value, either $y_j$ increases by $k_j > 0$ or $y_i$ decreases by $k_i > 0$ in the over-represented lie factor $L'_{j/i}$:

$$
\begin{aligned}
L'_{j/i} &= \frac{y'_i \downarrow - y'_j \uparrow}{y'_i \downarrow} \frac{x_j}{x_i - x_j} \\
&= \frac{(y_i - k_i) - (y_j + k_j)}{y_i - k_i} \frac{x_j}{x_i - x_j}
\end{aligned}
$$

We ask if the over-represented lie factor is greater than the well-represented lie factor $L_{j/i} = 1$:

$$
\begin{aligned}
L'_{j/i} &\overset{?}{>} L_{j/i} \\
\frac{(y_i - k_i) - (y_j + k_j)}{y_i - k_i} \frac{x_j}{x_i - x_j} &\overset{?}{>} \frac{y_i - y_j}{y_i} \frac{x_j}{x_i - x_j} \\
\frac{y_i - y_j - k_i - k_j}{y_i - k_i} &\overset{?}{>} \frac{y_i - y_j}{y_i} \\
y_i^2 - y_i y_j - k_i y_i - k_j y_i &\overset{?}{>} y_i^2 - k_i y_i - y_i y_j + y_j k_i \\
-k_j y_i &> y_j k_i \qquad \blacksquare
\end{aligned}
$$

Therefore, $L'_{j/i} > L_{j/i} = 1$.

**Case 3: Over-representation of $L'_{i/j}$, $x_j < x_i < 0$, $y_j < y_i < 0$.** Swap index such that $i \leftarrow j$ and $j \leftarrow i$. The proof is identical to case 2. Therefore, $L'_{i/j} > L_{i/j} = 1$.

**Case 4: Over-representation of $L'_{j/i}$, $x_j < x_i < 0$, $y_j < y_i < 0$.** Swap index such that $i \leftarrow j$ and $j \leftarrow i$. The proof is identical to case 1. Therefore, $L'_{j/i} > L_{j/i} = 1$.

**Case 5: Under-representation of $L'_{i/j}$, $x_i < x_j < 0$, $y_i < y_j < 0$.** To under-represent the true value, either $y_j$ decreases by $k_j > 0$ or $y_i$ increases by $k_i > 0$ in the under-represented lie factor $L'_{i/j}$:

$$\begin{aligned} L'_{i/j} &= \frac{y'_i \uparrow - y'_j \downarrow}{y'_j \downarrow} \frac{x_j}{x_i - x_j} \\ &= \frac{(y_i + k_i) - (y_j - k_j)}{y_j - k_j} \frac{x_j}{x_i - x_j} \end{aligned}$$

We ask if the under-represented lie factor is less than the well-represented lie factor $L_{i/j} = 1$:

$$\begin{aligned} L'_{i/j} &\overset{?}{<} L_{i/j} \\ \frac{(y_i + k_i) - (y_j - k_j)}{y_j - k_j} \frac{x_j}{x_i - x_j} &\overset{?}{<} \frac{y_i - y_j}{y_j} \frac{x_j}{x_i - x_j} \\ \frac{y_i - y_j + k_i + k_j}{y_j - k_j} &\overset{?}{<} \frac{y_i - y_j}{y_j} \\ y_i y_j - y_j^2 + k_i y_j + k_j y_j &\overset{?}{<} y_i y_j - y_j^2 - y_i k_j + y_j k_j \\ k_i y_j &< -y_i k_j \end{aligned} \quad \blacksquare$$

Therefore, $L'_{i/j} < L_{i/j} = 1$.

**Case 6: Under-representation of $L'_{j/i}$, $x_i < x_j < 0$, $y_i < y_j < 0$.** To under-represent the true value, either $y_j$ decreases by $k_j > 0$ or $y_i$ increases by $k_i > 0$ in the under-represented lie factor $L'_{j/i}$:

$$\begin{aligned} L'_{j/i} &= \frac{y'_i \uparrow - y'_j \downarrow}{y'_i \uparrow} \frac{x_j}{x_i - x_j} \\ &= \frac{(y_i + k_i) - (y_j - k_j)}{y_i + k_i} \frac{x_j}{x_i - x_j} \end{aligned}$$

We ask if the under-represented lie factor is less than the well-represented lie factor $L_{j/i} = 1$:

$$\begin{aligned} L'_{j/i} &\overset{?}{<} L_{j/i} \\ \frac{(y_i + k_i) - (y_j - k_j)}{y_i + k_i} \frac{x_j}{x_i - x_j} &\overset{?}{<} \frac{y_i - y_j}{y_i} \frac{x_j}{x_i - x_j} \\ \frac{y_i - y_j + k_i + k_j}{y_i + k_i} &\overset{?}{<} \frac{y_i - y_j}{y_i} \\ y_i^2 - y_i y_j + k_i y_i + k_j y_i &\overset{?}{<} y_i^2 + k_i y_i - y_i y_j - y_j k_i \\ k_j y_i &< -y_j k_i \end{aligned} \quad \blacksquare$$

Therefore, $L'_{j/i} < L_{j/i} = 1$.

**Case 7: Under-representation of $L'_{i/j}$, $x_j < x_i < 0$, $y_j < y_i < 0$.** Swap index such that $i \leftarrow j$ and $j \leftarrow i$. The proof is identical to case 6. Therefore, $L'_{i/j} < L_{i/j} = 1$.

**Case 8: Under-representation of $L'_{j/i}$, $x_j < x_i < 0$, $y_j < y_i < 0$.** Swap index such that $i \leftarrow j$ and $j \leftarrow i$. The proof is identical to case 5. Therefore, $L'_{j/i} < L_{j/i} = 1$.

##### 2.5.3 Derivation of lie factor of fold change for positive values

Consider $i$-$j$ combinations that satisfy $|x_i| > |x_j|$. Consider misrepresented but "well-behaved" graphs that (1) have positive values ($x_i > 0, x_j > 0$) and (2) do not visualize positive values with negative magnitude channels and vice versa ($\text{sgn}(x_i) = \text{sgn}(y_i)$, $\text{sgn}(x_j) = \text{sgn}(y_j)$).

**Case 1: Over-representation of $F'_{i/j}$, $x_i > x_j > 0$, $y_i > y_j > 0$.** To over-represent the true value, either $y_j$ decreases by $k_j > 0$ or $y_i$ increases by $k_i > 0$ in the over-represented lie factor $F'_{i/j}$:

$$\begin{aligned} F'_{i/j} &= \frac{y'_i \uparrow x_j}{y'_j \downarrow x_i} \\ &= \frac{y_i + k_i}{y_j - k_j} \frac{x_j}{x_i} \end{aligned}$$

We ask if the over-represented lie factor is greater than the well-represented lie factor $F_{i/j} = 1$:

$$\begin{aligned} F'_{i/j} &\overset{?}{>} F_{i/j} \\ \frac{y_i + k_i}{y_j - k_j} \frac{x_j}{x_i} &\overset{?}{>} \frac{y_i}{y_j} \frac{x_j}{x_i} \\ \frac{y_i + k_i}{y_j - k_j} &\overset{?}{>} \frac{y_i}{y_j} \\ y_i y_j + k_i y_j &\overset{?}{>} y_i y_j - k_j y_i \\ k_i y_j &> -k_j y_i \end{aligned} \qquad \blacksquare$$

Therefore, $F'_{i/j} > F_{i/j} = 1$.

**Case 2: Under-representation of $F'_{i/j}$, $x_i > x_j > 0$, $y_i > y_j > 0$.** To under-represent the true value, either $y_j$ increases by $k_j > 0$ or $y_i$ decreases by $k_i > 0$ in the under-represented lie factor $F'_{i/j}$:

$$\begin{aligned} F'_{i/j} &= \frac{y'_i \downarrow x_j}{y'_j \uparrow x_i} \\ &= \frac{y_i - k_i}{y_j + k_j} \frac{x_j}{x_i} \end{aligned}$$

We ask if the over-represented lie factor is less than the well-represented lie factor $F_{i/j} = 1$:

$$\begin{aligned} F'_{i/j} &\overset{?}{<} F_{i/j} \\ \frac{y_i - k_i}{y_j + k_j} \frac{x_j}{x_i} &\overset{?}{<} \frac{y_i}{y_j} \frac{x_j}{x_i} \\ \frac{y_i - k_i}{y_j + k_j} &\overset{?}{<} \frac{y_i}{y_j} \\ y_i y_j - k_i y_j &\overset{?}{<} y_i y_j + k_j y_i \\ -k_i y_j &< k_j y_i \end{aligned} \qquad \blacksquare$$

Therefore, $F'_{i/j} < F_{i/j} = 1$.

**Case 3: Over-representation of $F'_{j/i}$, $x_i > x_j > 0$, $y_i > y_j > 0$.** By Equation 2.41, $F'_{j/i}$ follows the opposite trend of case 1. Therefore, $F'_{j/i} < F_{j/i} = 1$.

**Case 4: Under-representation of $F'_{j/i}$, $x_i > x_j > 0$, $y_i > y_j > 0$.** By Equation 2.41, $F'_{j/i}$ follows the opposite trend of case 2. Therefore, $F'_{j/i} > F_{j/i} = 1$.

Now consider $i$-$j$ combinations that satisfy $|x_i| < |x_j|$.

**Case 5: Over-representation of $F'_{i/j}$, $x_j > x_i > 0$, $y_j > y_i > 0$.** Swap index such that $i \leftarrow j$ and $j \leftarrow i$. The proof is identical to case 3. Therefore, $F'_{i/j} < F_{i/j} = 1$.

**Case 6: Under-representation of $F'_{i/j}$, $x_j > x_i > 0$, $y_j > y_i > 0$.** Swap index such that $i \leftarrow j$ and $j \leftarrow i$. The proof is identical to case 4. Therefore, $F'_{i/j} > F_{i/j} = 1$.

**Case 7: Over-representation of $F'_{j/i}$, $x_j > x_i > 0$, $y_j > y_i > 0$.** By Equation 2.41, $F'_{j/i}$ follows the opposite trend of case 5. Therefore, $F'_{j/i} > F_{j/i} = 1$.

**Case 8: Under-representation of $F'_{j/i}$, $x_j > x_i > 0$, $y_j > y_i > 0$.** By Equation 2.41, $F'_{j/i}$ follows the opposite trend of case 6. Therefore, $F'_{j/i} < F_{j/i} = 1$.

The interdependence of the proofs suggests (1) the dependence relationship between $F_{i/j}$ and $F_{j/i}$ and (2) the necessity to define the sign of $i$-$j$ combinations.

##### 2.5.4 Derivation of lie factor of fold change for negative values

Consider $i$-$j$ combinations that satisfy $|x_i| > |x_j|$. Consider misrepresented but "well-behaved" graphs that (1) have negative values ($x_i < 0, x_j < 0$) and (2) do not visualize positive values with negative magnitude channels and vice versa ($\text{sgn}(x_i) = \text{sgn}(y_i)$, $\text{sgn}(x_j) = \text{sgn}(y_j)$).

**Case 1: Over-representation of $F'_{i/j}$, $x_i < x_j < 0$, $y_i < y_j < 0$.** To over-represent the true value, either $y_j$ increases by $k_j > 0$ or $y_i$ decreases by $k_i > 0$ in the over-represented lie factor $F'_{i/j}$:

$$\begin{aligned} F'_{i/j} &= \frac{y'_i \downarrow x_j}{y'_j \uparrow x_i} \\ &= \frac{y_i - k_i}{y_j + k_j}\frac{x_j}{x_i} \end{aligned}$$

We ask if the over-represented lie factor is greater than the well-represented lie factor $F_{i/j} = 1$:

$$\begin{aligned} F'_{i/j} &\overset{?}{>} F_{i/j} \\ \frac{y_i - k_i}{y_j + k_j}\frac{x_j}{x_i} &\overset{?}{>} \frac{y_i}{y_j}\frac{x_j}{x_i} \\ \frac{y_i - k_i}{y_j + k_j} &\overset{?}{>} \frac{y_i}{y_j} \\ y_i y_j - k_i y_j &\overset{?}{>} y_i y_j + k_j y_i \\ -k_i y_j &> k_j y_i \end{aligned} \qquad \blacksquare$$

Therefore, $F'_{i/j} > F_{i/j} = 1$.

**Case 2: Under-representation of $F'_{i/j}$, $x_i < x_j < 0$, $y_i < y_j < 0$.** To under-represent the true value, either $y_j$ decreases by $k_j > 0$ or $y_i$ increases by $k_i > 0$ in the under-represented lie factor $F'_{i/j}$:

$$\begin{aligned} F'_{i/j} &= \frac{y'_i \uparrow x_j}{y'_j \downarrow x_i} \\ &= \frac{y_i + k_i}{y_j - k_j}\frac{x_j}{x_i} \end{aligned}$$

We ask if the over-represented lie factor is less than the well-represented lie factor $F_{i/j} = 1$:

$$\begin{aligned}
F'_{i/j} &\stackrel{?}{<} F_{i/j} \\
\frac{y_i + k_i}{y_j - k_j}\frac{x_j}{x_i} &\stackrel{?}{<} \frac{y_i}{y_j}\frac{x_j}{x_i} \\
\frac{y_i + k_i}{y_j - k_j} &\stackrel{?}{<} \frac{y_i}{y_j} \\
y_i y_j + k_i y_j &\stackrel{?}{<} y_i y_j - k_j y_i \\
k_i y_j &< -k_j y_i
\end{aligned}$$

■

Therefore, $F'_{i/j} < F_{i/j} = 1$.

**Case 3: Over-representation of $F'_{j/i}$, $x_i < x_j < 0$, $y_i < y_j < 0$.** By Equation 2.41, $F'_{j/i}$ follows the opposite trend of case 1. Therefore, $F'_{j/i} < F_{j/i} = 1$.

**Case 4: Under-representation of $F'_{j/i}$, $x_i < x_j < 0$, $y_i < y_j < 0$.** By Equation 2.41, $F'_{j/i}$ follows the opposite trend of case 2. Therefore, $F'_{j/i} > F_{j/i} = 1$.

Now consider $i$-$j$ combinations that satisfy $|x_i| < |x_j|$.

**Case 5: Over-representation of $F'_{i/j}$, $x_j < x_i < 0$, $y_j < y_i < 0$.** Swap index such that $i \leftarrow j$ and $j \leftarrow i$. The proof is identical to case 3. Therefore, $F'_{i/j} < F_{i/j} = 1$.

**Case 6: Under-representation of $F'_{i/j}$, $x_j < x_i < 0$, $y_j < y_i < 0$.** Swap index such that $i \leftarrow j$ and $j \leftarrow i$. The proof is identical to case 4. Therefore, $F'_{i/j} > F_{i/j} = 1$.

**Case 7: Over-representation of $F'_{j/i}$, $x_j < x_i < 0$, $y_j < y_i < 0$.** By Equation 2.41, $F'_{j/i}$ follows the opposite trend of case 5. Therefore, $F'_{j/i} > F_{j/i} = 1$.

**Case 8: Under-representation of $F'_{j/i}$, $x_j < x_i < 0$, $y_j < y_i < 0$.** By Equation 2.41, $F'_{j/i}$ follows the opposite trend of case 6. Therefore, $F'_{j/i} < F_{j/i} = 1$.

The interdependence of the proofs again suggests (1) the dependence relationship between $F_{i/j}$ and $F_{j/i}$ and (2) the necessity to define the sign of $i$-$j$ combinations.

(8) Log Bar — Matplotlib 3.8.4 documentation.

(9) matplotlib.pyplot.bar — Matplotlib 3.9.2 documentation.

(10) Waskom, M. L. *Journal of Open Source Software* **2021**, *6*, 3021.

(11) Wickham, H., *ggplot2: Elegant Graphics for Data Analysis*, 2nd ed; Springer: New York, NY, 2016.

(12) MATLAB, 2024.

(13) Bar charts — geom_bar, en.

(14) GraphPad Prism.

(15) GraphPad Prism 10 User Guide - Advice: Bar graphs with log axes rarely make sense.

(16) I'm using a logarithmic scale for the Y axis of a bar graph. Prism plots numbers greater than 1 properly, but numbers below 1 as inverted bars. What can I do? - FAQ 219 - GraphPad.

(17) Graph tip - Don't use a log scale on a bar graph! - FAQ 1477 - GraphPad.
